# Partial epithelial-to-mesenchymal transition mediates profound gap closure through growth and fluidization

**DOI:** 10.64898/2026.07.09.737575

**Authors:** Han Jiang, Chaozhen Wei, Pengbo Wang, Jaivarsini Johnson, Nonthakorn Olaranont, Yifan Gu, Feiyang Chen, Jian Xu, Qi Wen, Min Wu, Yubing Sun

**Author notes:** Correspondence should be addressed to Yubing Sun and Min Wu. These authors contribute equally (Han Jiang, Chaozhen Wei).

## Abstract

Epithelial gap closure is essential for maintaining tissue integrity during development and wound healing. Previous studies have shown that closure of small gaps is driven by actomyosin purse-string contraction and traction forces generated at the gap edge. Here, we show that millimeter-scale circular gap closure in mouse epicardial (MEC1) monolayers is driven primarily by growth-mediated compressive stresses. Compared with MDCK monolayers, MEC1 cells close gaps more rapidly with reduced undulation near gap edge through coordinated tissue-wide extension-contraction. The collective closing dynamics can be modulated by partial epithelial–mesenchymal transition induction and Rho kinase inhibition. By integrating tissue and cell kinematic analyses, traction-force mapping, and a continuum framework that decomposes tissue strain rates into growth-, elastic-, and fluidity-related contributions, we reveal that growth-generated compression drives inward tissue flow, while elastic cell elongation and fluid-like tissue remodeling through cell–cell intercalation act synergistically to accommodate deformation and promote robust collective gap closure.

## INTRODUCTION

The migration of epithelial cells and fibroblasts is essential for the closure of gaps arising during development or following injury. Lamellipodia-mediated cell crawling is considered as the major migration mechanism for mesenchymal-like cells with nominal intercellular interactions^1,2^. For epithelial cells, pluricellular actin cables have been found in the wound edges in embryonic skin wound^3^ and intestinal epithelial cell lines^4^, leading to the theory that actin cables act as contractile ‘purse-strings’ to drive the gap closure^5^. Such purse-string mechanism has also been found in various development stages such as ventral enclosure in the *Caenorhabditis elegans*^6^ and the dorsal closure in *Drosophila* embryo^7^. It is widely accepted now that both cell crawling and ‘purse-string’ mechanisms contribute to epithelial gap closure, often using Madin-Darby Canine Kidney (MDCK) cells as an *in vitro* model system^8–12^. It is found that epithelial gap closure was initiated by cell crawling, and the late stage of closure was mediated by a combination of actin ring contraction and force transmission between the tissue and the substrates through focal adhesions^12^. Particularly, the epithelial cell crawling is a collective behavior that is often described using a leader-follower scheme: selected front cells exposed to void space become leader cells which generate significant traction forces through lamellipodia to drive the motion of the cell sheet. The specification of leader cells may be regulated by tugging forces generated by follower cells^13^, HGF/ERK signaling^14^, p53-p21 dependent cell cycle arrests^15^, and Notch1-Dll4 signaling^16^.

While initially driven by the active crawling forces of leader cells, high tissue fluidity, often characterized by cell-cell intercalation, has been found to facilitate the closure of small-scale gaps (i.e., around a few hundred micrometers in diameter and within cell doubling time)^17,18^. Cell-cell intercalation is also a critical component of a fundamental tissue remodeling motif, convergent extension, where tissues elongate along one axis while contracting laterally^19,20^. During convergent extension, cell-cell intercalation complements to cell elongation and alignment^21^ along the tissue elongation direction, and the resulting cell deformations are widely conceived as partially arising from elastic responses, as described in vertex models^22,23^. Notably, in the small-scale gap closure, the complementarity between cell elongation and cell-cell intercalations was found to be limited to the gap border^24^, and the tissue-scale extension-contraction characteristic of convergent extension has not been observed.

The current framework comprehensively describes the closure of small gaps of epithelial cells; however, it is insufficient to address many scenarios *in vivo* that involve larger gaps. The extended length scale, and correspondingly, time scale, make it essential to consider the nonnegligible effects of growth and cell state changes, such as epithelial-mesenchymal transition (EMT). For example, epicardial cells, a type of mesothelial cells that cover the outmost surface of the heart^25–27^, contribute to cardiac repair and regeneration in zebrafish and fetal mice^28,29^. For embryonic and adult epicardial cells, cell migration often occurs concomitantly with an EMT process in which cells *en route* retain a “partial EMT” phenotype^30–32^, as cells migrate as a cluster instead of individual cells. As cardiac injuries are often associated with millimeter scale gaps, it is critical to elucidate how tissue growth and (partial) EMT regulate profound gap closure.

In this work, we studied the cellular mechanisms involved in millimeter-scale gap closure of primary embryonic mouse epicardial (MEC1) cells. By performing tissue-scale imaging and traction force microscopy, we quantitatively characterized tissue growth, extension-contraction, and tissue-substrate mechanical interaction in profound gap closure. To elucidate the large-scale gap closure flow and deformation, we modeled the MEC1 monolayer as a thin plate with volumetric growth, elastic deformation, and fluidic rearrangements by extending the growth-elasticity theory^33^ to account for Maxwell-type viscoelasticity. Different from previous Maxwell viscoelasticity models with active force^34–36^ and their Stokes limit^37^, our fluidized growth-elasticity theory explained the observed tissue strain rate by growth-related, elastic-response-related, and fluidity-related strain rates through explicit rate and configuration decompositions. By connecting the model predictions and the cell and tissue shape dynamics from the experiments, our work reveals how growth, elasticity and fluidity are differentially utilized during growth-driven extension-contraction morphogenesis along the partial EMT spectrum.

## RESULTS

### MEC1 cells coordinately close profound gaps

To generate gaps with defined sizes and geometry and avoid artifact associated with scratching assays^8^, we used 3D-printed cylindrical stencils to generate circular gaps (diameter ≈ 1.7 mm) within confluent cell monolayers (**Supplementary Fig. 1** and **Methods**). We first examined the features of profound gap closure in epithelial cells using MDCK cells as a model (**Fig. 1a**, **Supplementary Video 1**). Consistent with prior findings in small gap closure of MDCK cells, both cell protrusions and supracellular actin bundles (“purse strings”) could be observed in the front cells (**Fig. 1a**, **Supplementary Video 1**). Also, we observed the emergence of several groups of leader cells and prominent undulation at the gap front, similar to what has been found in the re-epithelialization of keratinocyte to close profound circular wound^38^. In contrast, MEC1 cells closed the gap faster than MDCK cells (**Fig. 1a, b**) and showed reduced undulation at gap front, as quantified by the circularity of the remaining gap region (**Fig. 1c**, **Supplementary Fig. 2**). As epicardial cells undergo epithelial-to-mesenchymal transition (EMT) during cardiac development, we further evaluated whether epicardial gap closure was mediated by the EMT status of the cells.

**Figure 1.**
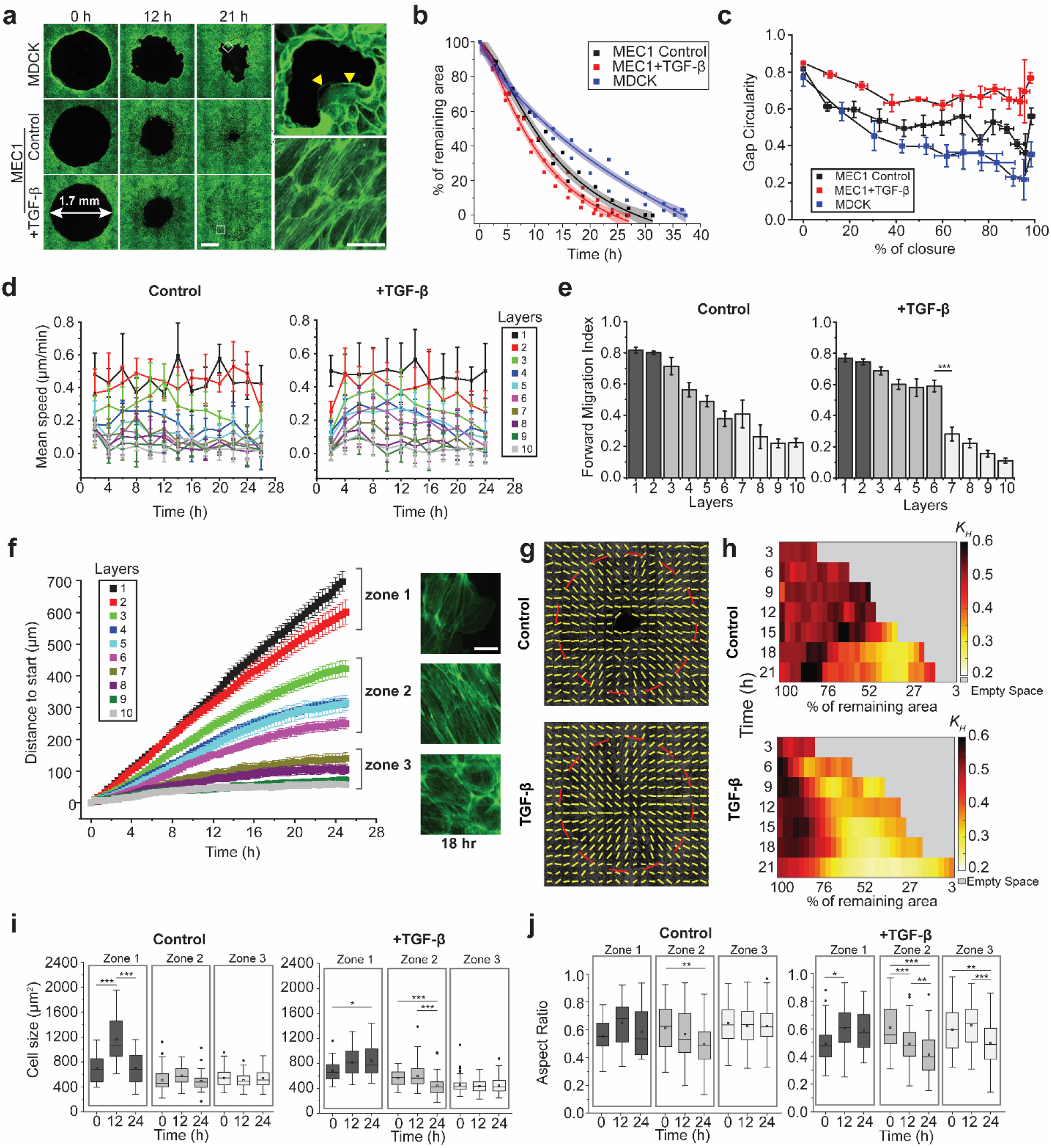
Cell migration kinematics and actomyosin remodeling during the gap closure process. (**a**) Left: Fluorescence images showing the actin staining of MDCK and MEC1 cells at 0, 12, and 21 h after stencil removal. Scale bar: 500 µm; Right top: Zoom-in images showing lamellipodia and actin cables in MDCK cells indicated by yellow arrows. Right bottom: Zoom-in images showing stress fibers in MEC1 cells. Scale bar for both images, 50 µm. (**b**) Plot showing the remaining gap area (normalized to initial gap area) decreases as a function of time. Solid lines show polynomial fitting and shaded areas show the 95% confidence intervals. *n* = 3 (**c**) Gap circularity changes as a function of percentage of gap closure. *n* = 3. The circularity is defined as 4π (area/perimeter^2^). (**d**) Mean speed of cells in each cell layer as a function of time. (**e**) Forward migration index (FMI) for cells in each layer. *FMI* = *x_i_*/ ∑ *d_i_*, where *x_i_* is the distance to start and *d_i_* is the distance to previous location. Only *P* values smaller than 0.05 between neighboring layers were indicated. (**f**) Plot showing the distance to the initial positions for TGF-β treated MEC1 cells in each cell layer as a function of time. Three zones are defined based on actin cytoskeletal structures at 18 h, as shown in the actin fluorescence images. Scale bar: 20 µm. (**g**) Actin fiber orientation analysis showing radial aligned actin fibers at 21 h for control and TGF-β treated MEC1 cells. (**h**) Colorimetric map showing *K_H_* values, which quantify the actin alignment, at different positions and time points. *n* = 4. (**i-j**) Average cell area (i) and cell aspect ratio (j) in each zone at 0, 12, and 24 h after stencil removal. Cell outlines were traced and analyzed using PAT-GEOM. Cell aspect ratio was defined as minor axis/major axis. *n* > 12 cells per cell layer were selected randomly and tracked over time. Data are represented as mean ± s.e.m. n.s., *P* > 0.05, *, *P* < 0.05, ***, *P* < 0.001.

By adding 5 μg/mL TGF-β1, we successfully induced EMT in MEC1 cells, supported by the expression of Slug and Vimentin (**Supplementary Fig. 3**). We found that TGF-β1 treatment slightly accelerated the gap closure speed and reduced the gap front undulation (**Fig. 1b, c**). Actin staining in Fig. 1a showed that unlike MDCK cells with predominately cortical actin, MEC1 cells possessed stress fibers while remaining connected with each other. Interestingly, we did not notice any “purse-string” like actin bundle formation in MEC1 cells throughout the entire process.

### Layer-dependent migration pattern in epicardial gap closure

To quantify spatiotemporal migration patterns of MEC1 cells, we analyzed the migration trajectories of the first ten cell layers, as cells further away showed negligible migration (**Fig. 1d-f**, **Supplementary Fig. 4, Supplementary Video 2, 3**). For both control and TGF-β1 treated conditions, the first two layers of cells move with a prominent speed compared to the trailing layers, which exhibited an initial speedup followed by a slowdown pattern (**Fig. 1d**). We also found that the forward migration index, measuring the directionality of the inward movement, decreased in the trailing layers (**Fig. 1e**). Interestingly, the TGF-β1 treated cells maintained a higher directionality in the 3-6^th^ layers, consistent with its overall higher circularity than the control condition (**Fig. 1c**). In addition, cells in the 3^rd^-6^th^ layers also had higher overall speed in the TGF-β1 treated condition (**Supplementary Fig. 4c).**

### EMT status regulates spatiotemporal evolution of actin stress fiber alignment

In TGF-β1 treated condition, we found cells in the 1^st^ and 2^nd^ layers (hereinafter referred to as zone 1) with the highest migration speed displaying prominent stress fibers without strong radial alignment (**Fig. 1f**). Cells in the 3^rd^-6^th^ layers (hereinafter referred to as zone 2) present strong radial alignment in their stress fibers associated with cell elongation. Cells in the 7^th^ – 10^th^ layers (hereinafter referred to as zone 3) with diminishing migration speed showed more prominent cortical actin enriched near cell boundaries, lacking any cell alignment. These observations were also confirmed by our actin fiber orientation analysis (**Fig. 1g**). We further quantitatively investigated spatiotemporal evolution of such stress fiber orientation. Considering a distribution of actin fiber orientation with *ρ*(*α*), the local orientation density function, and 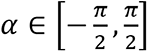 the angle between the actin and radial directions, we utilized a structure parameter, 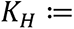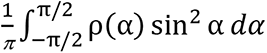, to quantify the distribution of tissue-level actin fiber orientation^39^ (see **Methods**). By definition, the actin fiber distribution is more radially (circumferentially) aligned as *K_H_* decreases (increases) and *K_H_* = 0.5 indicates that the actin fiber distribution is not aligned with either direction when *ρ*(*α*) ≡ 1. As shown in **Fig. 1h and Supplementary Fig. 5a**, both control and treated conditions have the radial alignment formulating during the closure, with middle regions showing the strong radial alignment. Notably, TGF-β1 treatment enhanced the overall radial alignment in both the initiation time (6 h vs. 18 h) and spatial coverage. MEC1 cells consistently showed a lower *K_H_* value compared to MDCK cells, which has a *K_H_* value around 0.5 (**Supplementary Fig. 5b**).

### Cell size and shape changes during gap closure

Consistent with prior reports in small gap closure, MDCK cells maintain epithelial phenotypes and cell shape throughout the gap closure process with transient cell shape changes in leader cells^13^. In contrast, the front layers (zone 1) of untreated MEC1 cells showed a drastic increase in cell areas in the midst of gap closure and reduced to ground level towards the end of the gap closure (**Fig. 1i**). TGF-β1 treated MEC1 cells showed modest increase in cell areas in zone 1 but a significant cell area reduction in zone 2 at the late stage (**Fig. 1i**). In both control and TGF-β1 treated conditions, front cells were consistently larger than rear cells (**Fig. 1i**). The aspect ratio analysis further confirmed that TGF-β1 treatment promoted cell elongation in zone 2 and to a less extent, zone 3 (**Fig. 1j**). These observations suggest spatiotemporal heterogeneous behavior of MEC1 cells during migration.

### The effects of initial gap curvature on migration patterns

We next ask whether the gap curvature mediates the differential cell migration patterns. Circular gaps with diameters of 1 mm and 300 µm (**Fig. 2a, d**), and unilateral straight gaps (**Fig. 2a, g**) were generated, and MEC1 cells were seeded and treated with TGF-β1, representing a range of curvatures from approximately 0 to 6.67 mm^-1^. For the larger curvatures, we found that cells prone to elongation and radial alignment (i.e., zone 2 in Fig. 1f) started to diminish with decreasing gap sizes and only layer 1 cells displayed a high migration speed in both conditions (**Fig. 2a-f**). Notably, the 2^nd^ layer had about half of the migration speed as the 1^st^ layer in the 1 mm gap, while only having one third of the migration speed in the 300 µm gap. The migration speed of the front cells in 1 mm gaps was comparable with 1.7 mm gaps (∼ 0.5 µm/min), which was significantly higher than the cells in 300 µm gaps condition (*p* < 0.01). Radial alignment in cells or stress fibers was not observed in 300 µm gaps condition.

**Figure 2.**
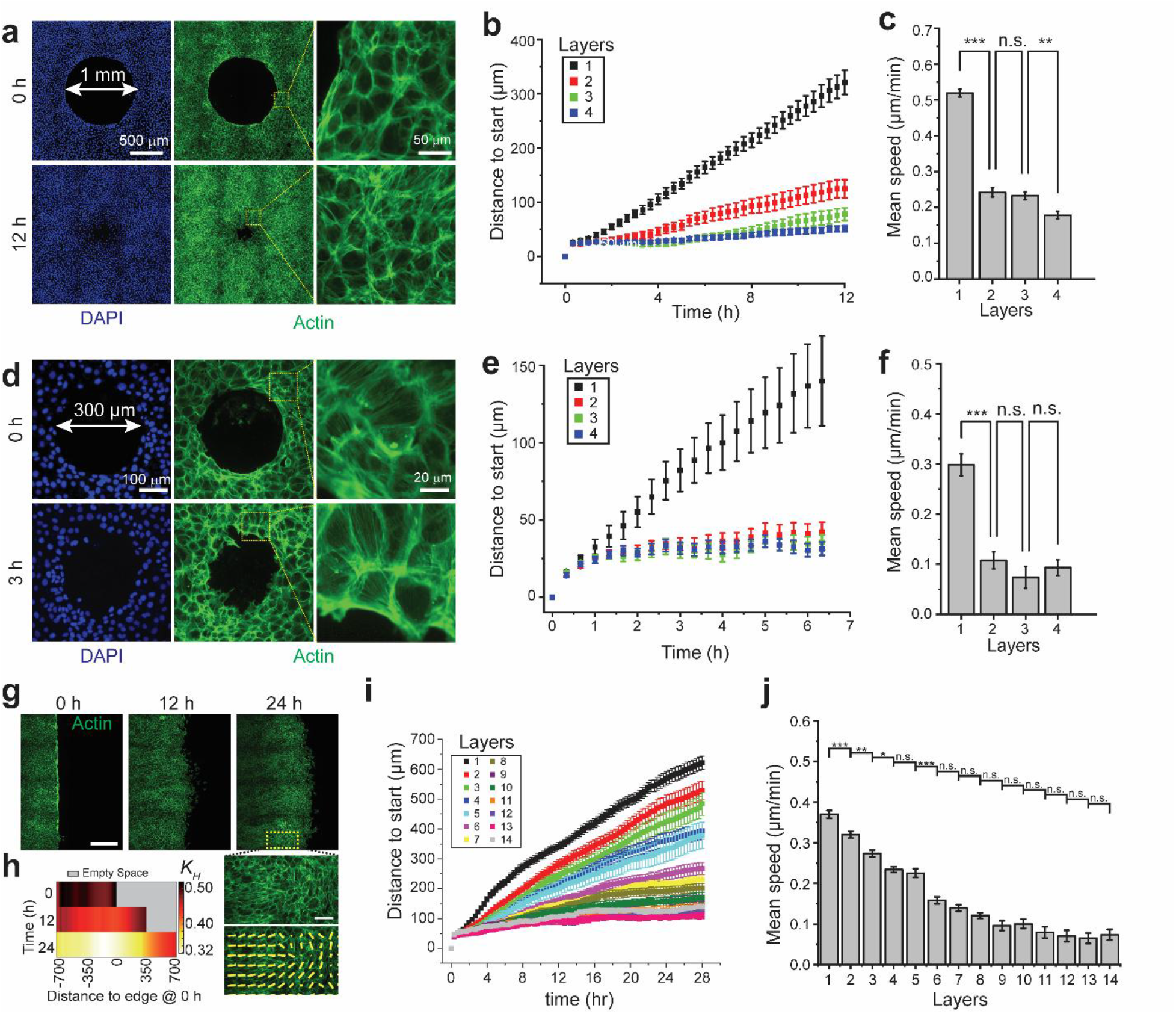
Gap size and geometry regulate gap closure patterns. (**a**) Representative fluorescence images showing actin and cell nuclei for gaps with diameters of 1 mm. (**b**) Plot showing the distance to the initial positions for cells in the first four layers as a function of time. The width of each layer was defined as 49 µm. *n* = 3. (**c**) The mean speed of cells within each layer. (**d-f**) Images and plots showing gap closure dynamics in gaps with diameters of 300 µm. The width of each layer was defined as 24.5 µm. Scale bar in **d**, 100 µm. *n* = 3. (**g**) Top: Representative fluorescence images showing actin staining of cells closing a straight unilateral gap and bright field image showing the cell tracking. Scale bar, 500 µm. Bottom: Zoom-in images and fiber orientation analysis showing cells with actin alignment perpendicular to the wound boundary. Scale bar, 200 µm. (**h**) Colorimetric map showing *K_H_* values at different positions and time points. *K_H_* was defined to be 0 when fibers were perpendicular to the wound boundary, and 1 when they are in parallel. *n* = 3. (**i-j**) Plots showing the distance to the initial positions and mean speed for cells in each layer. *n* > 9 cells in each layer in each sample and three independent experiments were performed. All the results were obtained using MEC1 cells treated with TGF-β1. Data are represented as mean ± s.e.m. n.s., *P* > 0.05, *, *P* < 0.05, **, *P* < 0.01, ***, *P* < 0.001.

For the unilateral straight gaps, the front cells traveled with a slower speed than the 1.7 mm circular gap, while the active migration region extended to a longer range. Cell migration speed dropped more gradually from the front for straight gaps compared with the 1.7 mm circular gaps. The mean speed dropped to half of the front cell speed in the 5^th^-6^th^ layers, compared to the 3^rd^ and 4^th^ layers for 1.7 mm circular gaps (**Fig. 2i, j**). We also found that MEC1 cells gradually reorientated to perpendicular to the gap boundary direction, particularly about 350 µm behind the front layer (**Fig. 2h**). To summarize, while there was not a clear correlation between front cell migration speed and gap curvature, the collective migration extended to a longer range for larger gaps with smaller curvature.

### Traction stress maps during MEC1 gap closure

The lack of supracellular contractile actin rings excludes the possibility of purse string as the driving force of MEC1 gap closure. Thus, we examined whether the crawling of the front cells generated large contraction forces to drive the gap closure. Previous studies have found an increase of p53/p21 expression in the leader cells during collective migration of MDCK cells^15^, leading to the cell cycle arrest and multinucleated cells that drive the migration of cell clusters. Similarly, large, multinucleated cells were also found in zebrafish epicardium in the wound front, which migrated at a high velocity and generate large mechanical tensions^40^. Thus, we first examined whether leader cells also existed and drove the profound gap closure in MEC1 cells. As shown in **Fig. 3a, b**, p53+ cells distributed uniformly across the monolayer and no major upregulation of p53 was found in the front cells. In addition, although some multinucleated cells were observed (**Supplementary Video 2, 3**), those cells were found randomly located and showed reduced migration speed compared to their neighboring cells.

**Figure 3.**
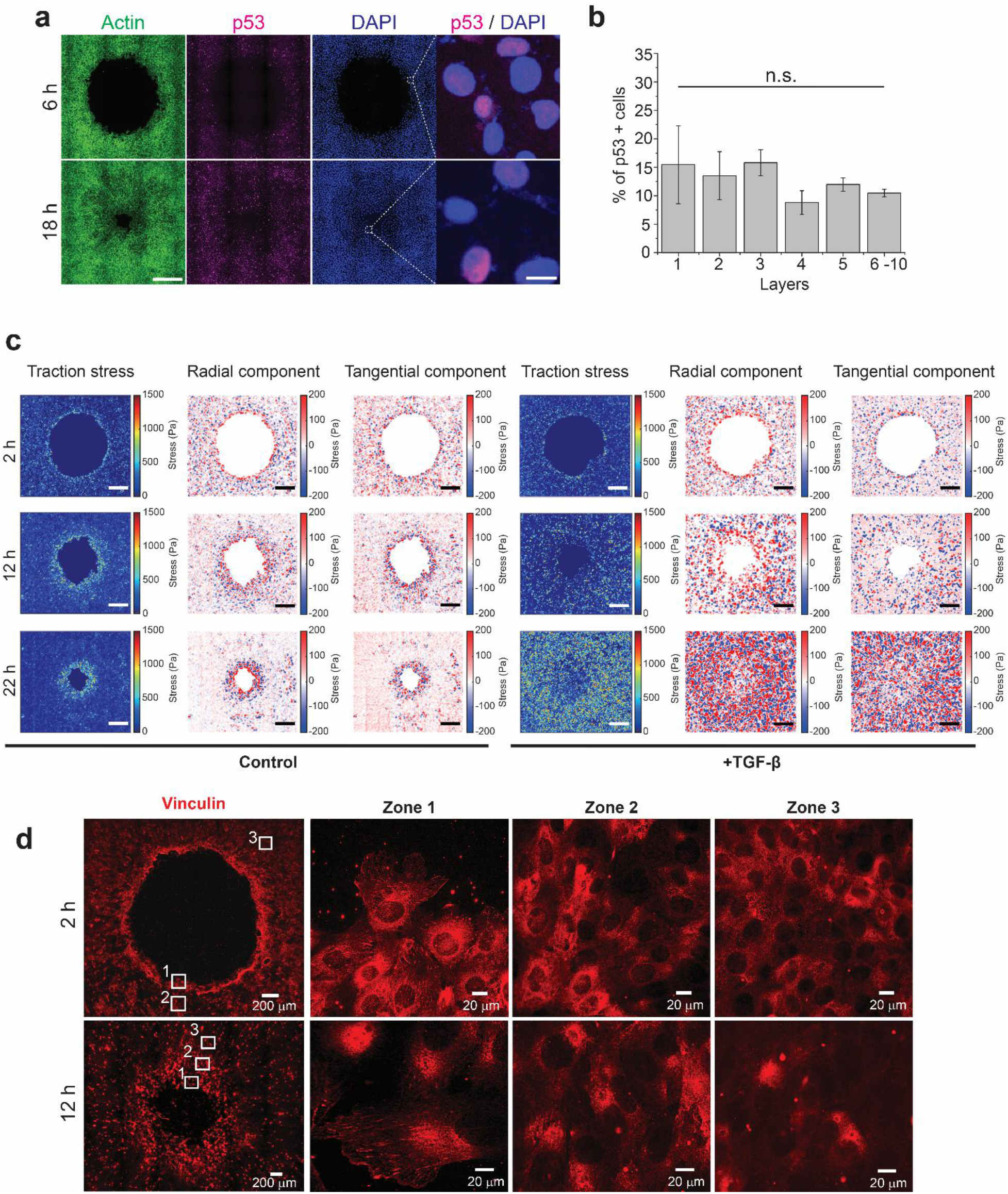
Traction stress maps during MEC1 gap closure. (**a**) Representative fluorescence images showing the actin and p53 staining in TGF-β1 treated MEC1 cells at 6 and 18 h. Scale bar: 500 μm for the first three panels and 20 μm for the last panel; *n* = 3 (**b**) Bar plots showing the percentage of p53 positive cells in each layer at 18 h. No statistical significances were found using one-way ANOVA analysis. (**c**) Heatmaps showing the magnitude, tangential component, and radial component of traction stress in control and TGF-*β*1 treated MEC1 cells at 0, 12, and 22 h. Scale bar: 500 μm. (**d**) Fluorescence images showing vinculin expression in MEC1 cells.

To further clarify the role of front cells, we performed traction force microscopy for MEC1 gap closure process. Traction force mapping for a large area possesses a significant challenge due to inevitable imaging distortion and misalignment during image stitching. We combined an improved fluorescence microbeads-based traction force microscopy with a customized image correction algorithm and successfully obtained traction force maps for the entire gap region (see Methods for details). We found that in the control condition, cell traction forces concentrated in the front layers uniformly, suggesting the front cells pulled the cell sheet collectively (**Fig. 3c**), in contrast to sporadic leader cell drive clusters found in MDCK gap closure^12^. TGF-β1 treatment increased the uniformity of traction stress distribution in expanded areas. We also noticed interesting evolution of the traction maps. At the early stage, the traction stress was most significant at the front along the radial direction in both conditions (2 h, **Fig. 3c**). At later stages of the control group, the region with significant traction stress expanded locally (12 h and 22 h, **Fig. 3c** and **Supplementary Fig. 6**), where the traction stress in the following layers did not show clear directionality. In contrast, the traction stress in the TGF-β1 treated group showed significant radial directionality in expanded areas but reduced total traction stress in front cells (12 h, **Fig. 3c** and **Supplementary Fig. 6**). After closure, the traction stress remained significant in the treated TGF-β1 group while directionality disappeared (22 h, **Fig. 3c** and **Supplementary Fig. 6**).

We further examine the cell-substrate interactions by staining vinculin at early and late stages of gap closure (**Fig. 3d**). We found that vinculin expressions were elevated in the front 1-2 layers of cells at 2 h and expanded to the next 3-5 layers at 12 h. The weak vinculin expression in the rear was consistent with the reduced traction forces. Interestingly, the increased traction force and adhesion signals in the front cells did not lead to YAP-dependent mechanical activation^41^, which was also not influenced by the EMT status (**Supplementary Fig. 7**). This is in contrast with YAP-dependent leader cell selection found in other systems^42,43^.

### Cell proliferation regulates profound gap closure

A main feature of profound gap closure is the extended gap closure time, during which cell proliferation cannot be neglected. Our traction force measurement suggested that the pulling forces generated by the front cells were localized and might not be sufficient to drive forward movement of the monolayer. We thus hypothesize that cell proliferation leads to compressive forces within the cell monolayer, which in turn push forward the following cells. To test whether MEC1 cells underwent active mitosis, we first stained phosphorylated histone H3 (pHH3), which marks mitotically active cells in late G2/M phases, at different time points (**Fig. 4a**). We found that throughout the full gap closure process, MEC1 cells maintained at a stable proliferation rate of ∼0.022 hr^-1^, calculated based on mitosis time we measured using live-cell microscopy (**Fig. 4b**, see **Methods**)^44^. Spatially, we also observed that the cell proliferation appeared uniformly within the monolayer (**Fig. 4c**), suggesting various mechanical status did not significantly influence the cell proliferation. To evaluate the role of cell proliferation, we treated cells with aphidicolin^45^. We found that aphidicolin effectively reduced cell proliferation rate throughout the entire gap closure process (**Fig. 4d, e**, **Supplementary Fig. 8**). We tracked the cell migration in the first 5 layers of cells and found that aphidicolin treated MEC1 cells had a reduced gap closure speed (∼28 h for a full closure compared with 21 h in only TGF-β1 treated group, **Fig. 4f**). At individual cell level, only the first layer maintained a distinctively high velocity, which also decreased overtime (**Fig. 4g**). This is in contrast with the TGF-β1 only condition where the migration speed of the zone 1 cells only started to decrease close to the end **(Fig. 1e**). The cell migration speed drastically reduced starting from layer 2 (**Fig. 4h**). Such discrepancy in migration speed led to disruption in cell-cell contacts and cell area increase in the gap front, while actin alignment was still observed in the rear (**Fig. 4d**). Together, these data suggest that cell proliferation is critical to the migration pattern of the gap closure. However, the cell and fiber alignment seemed to still form with tissue flow towards the gap center.

**Figure 4.**
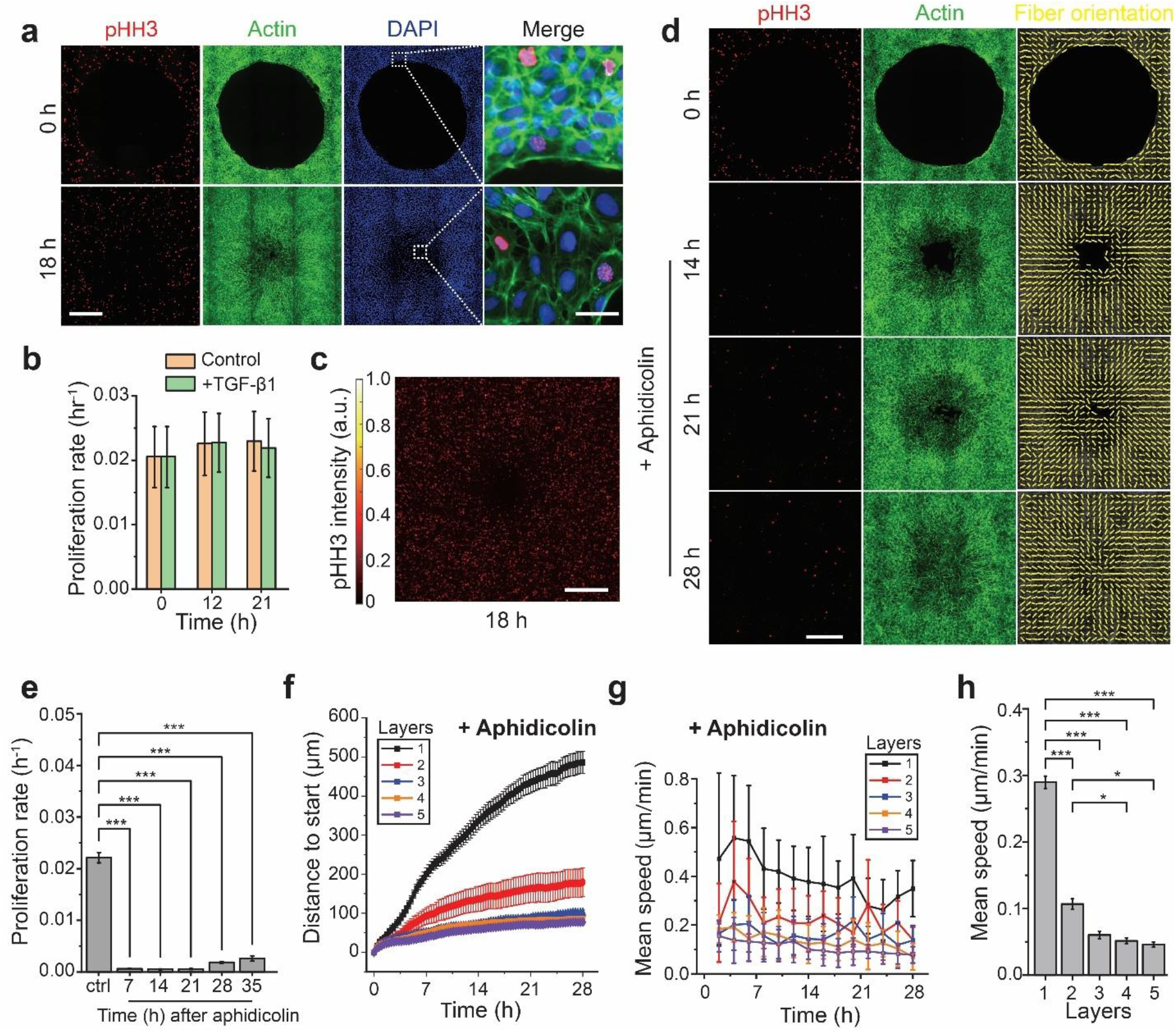
Cell proliferation regulates MEC1 gap closure. (**a**) Representative fluorescence images showing the phosphorylated histone H3 (pHH3) and actin staining in TGF-β1 treated MEC1 cells at 0 and 18 h. Scale bar: 500 μm for the first three panels and 50 μm for the last panel; Cells were counterstained with DAPI. (b) Bar plots showing proliferation rate of MEC1 at different time points. (**c**) Colorimetric map showing averaged pHH3 distribution at 18 h. *n* = 17 (**d**) Representative fluorescence images showing the phosphorylated histone H3 (pHH3) and actin staining and orientation analysis in MEC1 cells treated with both TGF-β1 and aphidicolin at 0, 14, 21, and 28 h. Scale bar: 500 μm. (**e**) Bar plots showing the proliferation rate with and without aphidicolin treatment. (**f**) Plot showing the distance to the initial positions for TGF-β1 and aphidicolin treated MEC1 cells in the first five layers as a function of time. (**g**) Plot showing the mean speed of aphidicolin treated cells in each cell layer as a function of time. (**h**) Bar plot showing the mean speed for aphidicolin treated cells in each layer. Data are represented as mean ± s.e.m., *, *P* < 0.05, ***, *P* < 0.001.

### Tissue growth and extension-contraction are critical in MEC1 gap closure

Given the traction map and proliferation analysis, we hypothesized that the profound gap closure in MEC1 monolayer underwent tissue area growth and extension-contraction deformation similar to convergent extension. We obtained strain rates 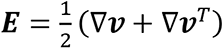, where ***v*** is the velocity, at different time points from particle image velocimetry (see Methods) and extracted the distribution of the spectral radius of 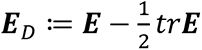 that measured the rate of extension-contraction, and *trE* (= *div**v***) that measured the area growth rate, both in control and TGF-β1 treated conditions (**Supplementary Fig. 9**). In both cases, we found the area growth rate increased and then decreased along the distance from the gap border at early and middle stages (4 and 12 h). Strikingly, the front region showed area contractions close to the end (24 h). In both control and TGF-β1 treated conditions, the extension-contraction rate stayed positive in time, and the direction of extension aligned mostly with the local radial axis (**Supplementary Fig. 9**).

These measurements are corroborated by front and rear tissue patch deformations (**Fig. 5a**, see **Methods** for details). The shape changes of a quadrilateral-shaped tissue were tracked based on the migration trajectory of cells at four vertices (**Fig. 5b**). We found that tissue patches in the gap front in the control condition showed a significant growth and extension-contraction characterized by increases in aspect ratio, area, and radial length, and decrease in circumferential length, while the tissue patches in the rear remained almost unchanged in aspect ratio (**Fig. 5c**, control condition). In contrast, the rear tissue patches in the TGF-β1 treated condition showed moderate extension-contraction (**Fig. 5c**, TGF-β1 treated condition). This difference coincided with a higher migration speed and directionality for the rear layers in the TGF-β1 treated condition (**Fig. 1e, f**). Last, the late-stage contraction from the PIV analysis was supported by the decrease of area in front patches close to 24 hours in both control and treated conditions (**Fig. 5c**).

**Figure 5.**
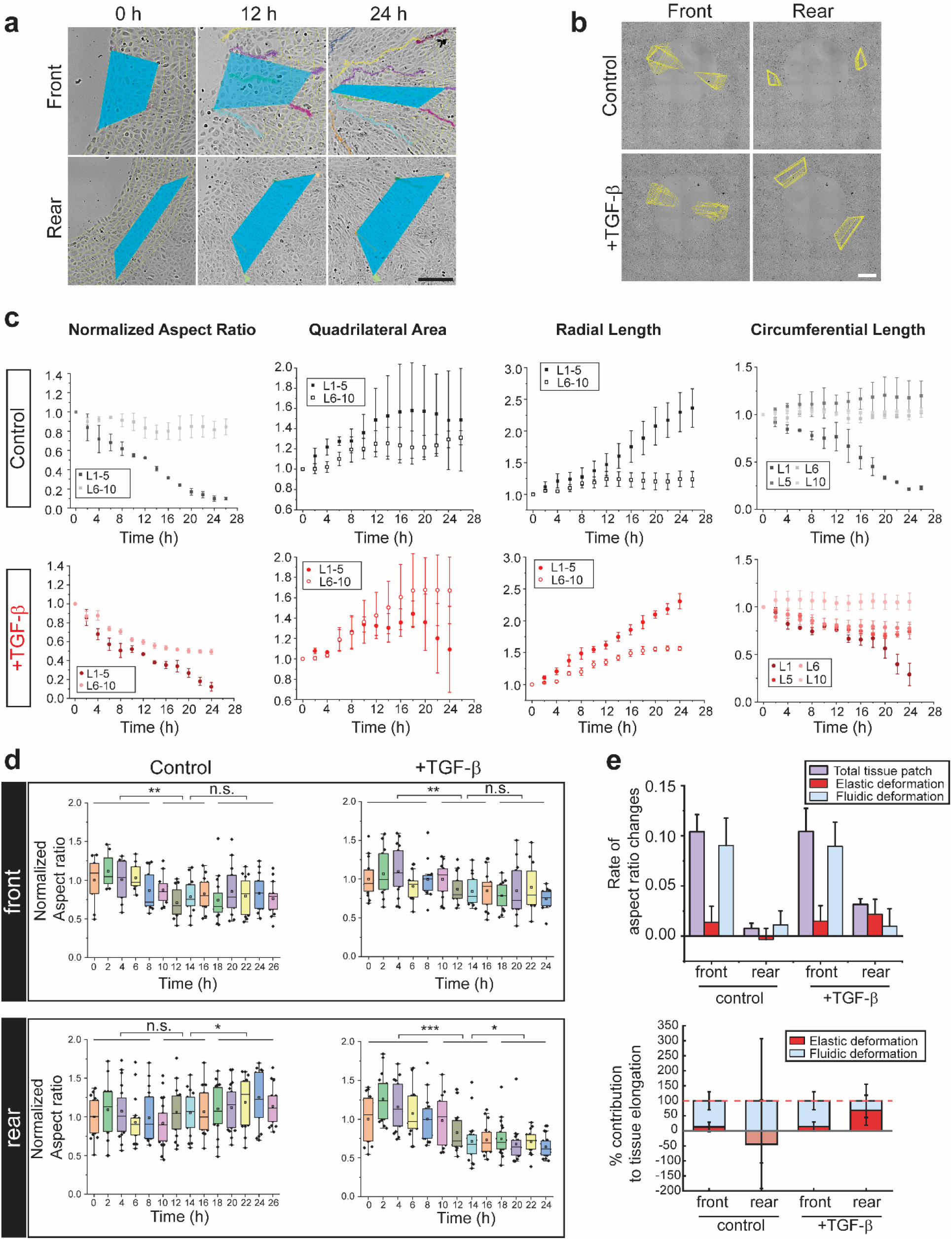
Tissue fluidity facilitates gap closure. (**a**) Representative brightfield images showing examples of tissue patches. Blue shaded areas indicated the quadrilateral shaped tissue patches and lines indicated sscell migration trajectories. Scale bar, 400 µm (**b**) Brightfield images showing time-lapse tracking of quadrilateral shaped tissue patches throughout the gap closure process. Scale bar, 400 µm. (**c**) Quantification of tissue patch deformation. Plots showing the tissue patch deformations under control or TGF-β1 treatment conditions. For each tissue patch (quadrilateral), the length connecting cells in the same layer is defined as the circumferential length, and the length connecting cells in different layers is defined as the radial length. The aspect ratio was determined by the ratio of the shorter radial length divided by the average circumferential length. (**d**) Cell aspect ratio changes throughout the gap closure process. Aspect ratio of cells at different timepoints, defined by the ratio of short axis to the long axis, was determined by ImageJ using cell outlines. The values were then normalized to the aspect ratio at 0 hour. Cells in the front and rear tissue patches were analyzed respectively. *n* > 11 cells for each time point. (**e**) Analysis of mean rate of the change of the aspect ratio of the total tissue patches, and relative contributions from cell elongation and intercalation, respectively. See **Methods** for details. Data are represented as mean ± s.e.m. n.s., *P* > 0.05, *, *P* < 0.05, **, *P* < 0.01, ***, *P* < 0.001.

### Cell elongation and intercalation are responsible for extension-contraction

The tissue-level extension-contraction can be due to cell elongation, cell-cell intercalation^17^, and oriented cell division (OCD)^46^. To test whether OCD caused the convergent extension, we first measured the nucleus-to-Golgi polarity by staining Golgi apparatus^47^, and did not notice any global polarity (**Supplementary Fig. 10a**). We further quantified the cell division angle by tracking dividing cells (**Supplementary Fig. 10b-c)**. Directional cell division was observed in neither front nor rear cells. Thus, we concluded OCD was not a cause of the convergent extension.

We then compared the tissue-patch and cell aspect ratio changes and quantified the tissue extension-contraction related to cell elongation and that related to intercalation (**Fig. 5e**). For both control and TGF-β1 treated conditions, cell intercalation contributed to most of the tissue extension-contraction in the front region. In the rear region, the control cells were compressed slightly in the radial direction, while a significant cell elongation was found in the TGF-β1 treated condition, dominating the tissue deformation, resembling more of an elastic response to local mechanics rather than a fluidic response.

### Mechanobiological modeling decomposes the inward flow and extension-contraction

Our experimental results so far revealed that the inward flow patterns and extension-contraction were associated with cell growth/proliferation, cell elongation, and cell-cell intercalation. Between the control and the TGF-β1 treated condition, the spatiotemporal patterning of these activities varied except for the cell proliferation. Thus, we assumed that the inward flows were primarily induced by growth at all times in both conditions and were mediated differently mechanically. To elucidate the mechanical difference between the two conditions, we have developed a continuum theory of soft tissue flow and deformation in the geometry of a circular gap on a thin plate^48^.

At each instant, the tissue was described as a three-dimensional incompressible fiber-reinforced neo-Hookean material, with reinforcing fibers restricted in the two-dimensional plane, representing the actin network. Consistent with the experiment, we modeled the initial distribution of the fiber orientation by the initial structure parameter 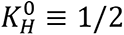 (**Supplementary Fig. 5**) and reasoned that the evolving fiber orientation 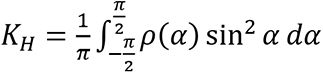 was partially due to the local elastic deformation that varied the density function *ρ*(*α*) (see **Methods** for details). Tissue growth due to cell growth and division generated elastic stresses and deformations within the monolayer, where the fluidity arose from stress-driven local mass rearrangements as in Maxwell models^35,49^. The model generated inward flow patterns and associated deformations with four free model parameters: growth rate γ, planar rearrangement tendency β, related to the cell-cell intercalation, third-dimensional thickness rearrangement ς, and level of planar fiber reinforcement η relative to the ground neo-Hookean material. Although all model parameters may vary spatiotemporally, we assumed them to be constant within each experimental condition (control versus TGF-β1 treated) for simplicity. See ref. 48 for the model details and summary of other modeling approaches.

We fitted our model to cell-layer-specific velocity data from both control and TGF-β1 treated conditions, and the spatiotemporal dynamics of the velocity agreed well with the experiment, including an early speed up followed by a later slow down^48^. The TGF-β1 treated condition presented a higher speed than the control conditions in the trailing layers (**Fig. 6a**), and an early increase and later-stage contraction (slowdown of increase) in the area for the front (rear) patches (**Fig. 6b**). Both characteristics are consistent with the experiments (**Fig. 1e**, **Supplementary Fig. 4c** and **Supplementary Fig. 9**). In addition, the front patches in both simulated control and treated conditions showed drastic decrease of aspect ratio (**Fig. 6c**).

**Figure 6.**
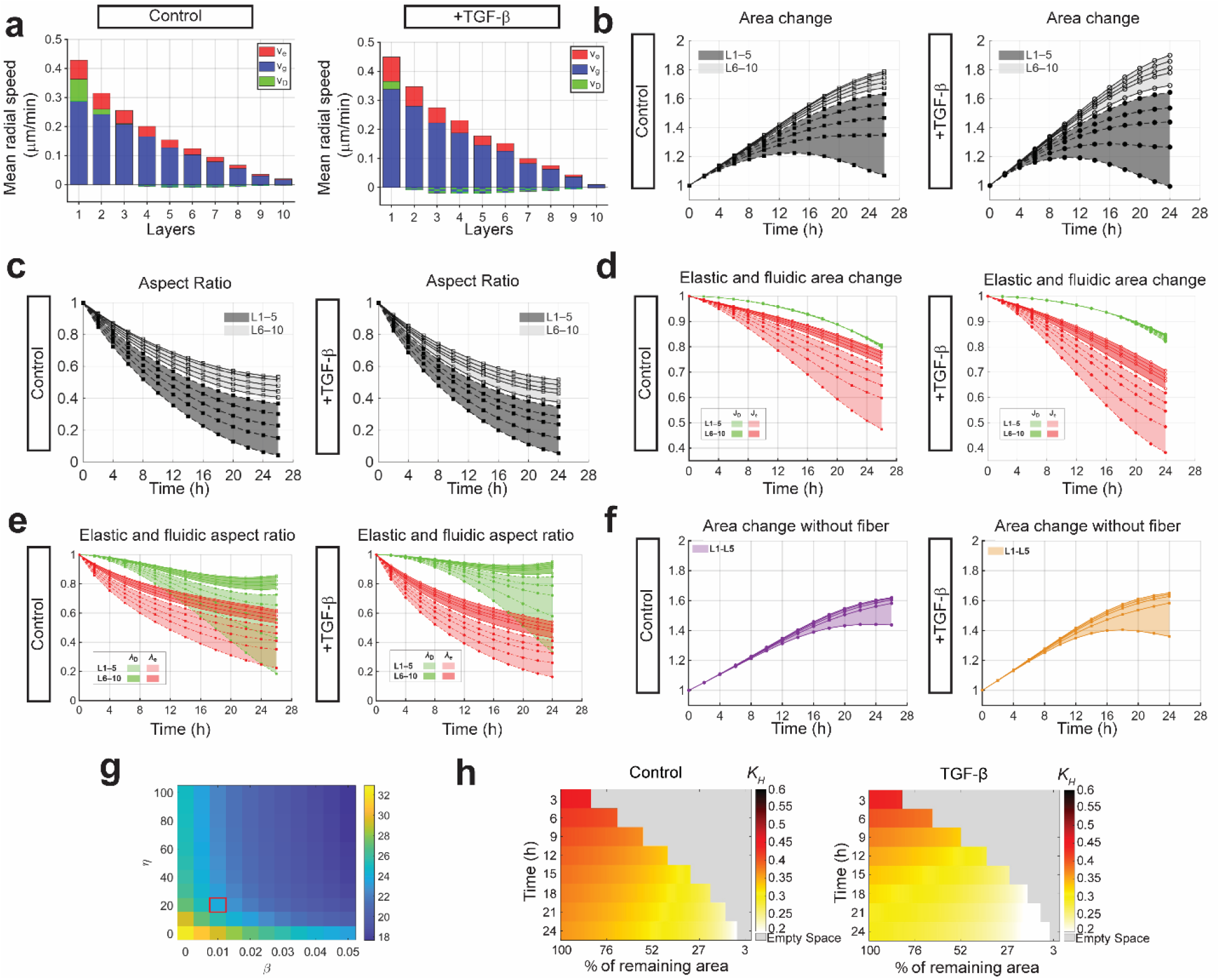
Model fitting and decomposition of the inward flow and deformation. (**a**) Mean speed of the 10 layers from the best-fit, with components due to growth (v_g_), elastic deformation (v_e_), and fluidic rearrangement (v_D_) predicted by the model. (**b, c**) Plots showing the tissue patch area and aspect ratio vs. time in the 5 front layers (L1-5) vs the 5 rear layers (L6-10), respectively. (**d**) Plot showing the change of area due to elastic deformation (J_e_) and fluidic rearrangement (J_D_) predicted by the model. (**e**) Plot showing the change of aspect ratio due to elastic deformation (λ_e_) and fluidic rearrangement (λ_D_) predicted by the model. (**f**) Plots showing the tissue patch area vs. time in the 5 front layers (L1-5) from the best-fit simulation when fiber-reinforcement was turned off (η = 0, both control and treated). (**g**) Prediction of total time (hr) needed to reach a gap radius of 200 *μm* when fiber-reinforcement level η and planar rearrangement tendency β vary for the TGF-β1 treated condition (marked in red square). (**h**) Colorimetric map showing predicted *K_H_* values, which quantify the actin alignment from the simulation, at different positions and time points for simulated control and TGF-β1 treated conditions.

We decomposed the simulated deformation into growth, elastic, and fluidic components (see **Methods**). While growth continued to have a significant contribution on both front and rear area changes, the decomposition showed the late-stage contraction at the front was mainly due to planar elastic compression (**Fig. 6d**, L1-5, Je). For the aspect ratio, the decomposition revealed that the elastic deformation contributed to the aspect ratio change at all times (red curves in **Fig. 6e**), while the fluidic rearrangement only began to contribute to the aspect ratio change at later stages (green curves in **Fig. 6e**). In addition, the TGF-β1 treated condition indeed had a higher elastic component than the control counterpart in the rear regions (L6-10 red curves, **Fig. 6e**), corroborating the cell elongation observed in the rear regions only for the TGF-β1 treated condition (**Fig. 5d**). We further decomposed the inward flow into the three corresponding components, and the more localized front layer speed in the control condition was explained by a higher level of fluidity than the TGF-β1 treated condition (**Fig. 6a**).

While we did not identify significant fiber-associated difference among the two conditions from the model, we found that the fiber-reinforcement was crucial for the area contraction to happen. When we fitted the model to velocity data by removing the fiber-reinforcement (η=0), the simulated front tissue patch completely lost the late-stage contraction in the control condition, while in the TGF-β1 treated condition only a slight late-stage contraction remained, limited to the first cell layer (**Fig. 6f**). Furthermore, our parameter study showed that both fiber reinforcement and planar rearrangement facilitate faster tissue closure; however, near the TGF-β1 treated condition, closure speed is more sensitive to changes in planar rearrangement than to changes in fiber reinforcement (**Fig. 6g**).

Lastly, we found that the enhanced radial fiber alignment in the TGF-β1 treated condition could be partially explained by the model (**Fig. 1h** and **Fig. 6h**). Although the evolution of the fiber orientation *K_H_*was modeled solely as a consequence of local elastic deformation, we reason that the TGF-β1-dependent radial fiber alignment is likely driven by actin remodeling^50^, guided by the directionality cues provided by increased elastic deformation under TGF-β1 treatment. Specifically, the larger elastic deformation inferred in the TGF-β1 treated condition may provide stronger directional cues, thereby contributing to the enhanced radial alignment. For more model features and theory results, see ref. 48.

### Enhanced tissue fluidization facilitates efficient gap closure

Our continuum theory predicted that enhanced planar rearrangement could speed up the gap closure significantly under varying fiber-reinforcement conditions. To validate the prediction, we treated cells with both TGF-β1 and a Rho kinase inhibitor (Y27632, or RI), which has been shown to enhance tissue fluidity^17^. Indeed, we found that RI treatment significantly shortened the gap closure time from 21 h to 16 h (**Fig. 7a, b**). Cell kinematic analysis showed that RI treatment led to an increased cell speed in the front two layers (zone 1) and expanded the zone 2 to five layers (**Supplementary Fig. 11a**). The cells in zone 2 also showed larger forward migration index compared with TGF-β1 only condition (**Supplementary Fig. 11b**). The accelerated cell migration was not due to changes in cell proliferation, which remained at a constant level (**Supplementary Fig. 11c**). Our model-specific velocity decomposition analysis inferred a globalized tissue fluidity in comparison with the control and TGF-β1 treated conditions (**Fig. 7c** and **Fig. 6a**). We also noticed a reduced circularity of gap in RI treated cells, which might result from increased cell-cell intercalations (**Fig. 7d**). To confirm that RI enhanced tissue fluidity to promote gap closure, we measured cell elongation (**Fig. 7e**) and tissue patch deformation (**Fig. 7f**) and calculated the relative contribution from cell elongation and intercalation (**Fig. 7g-h**). We found a lack of cell elongation both in the front and rear regions (**Fig. 7d**), and the tissue strain analysis also demonstrated the dominating role of tissue intercalation in both front and rear cells (**Fig. 7g-h**). Traction force microscopy revealed that RI treatment reduced the global cell traction and disrupted the radial orientation at gap front (**Fig. 7d**). The reduced traction and increased gap-closure speed in RI-treated cells provided further evidence that traction forces were not the leading driving force for the MEC1 gap closure.

**Figure 7.**
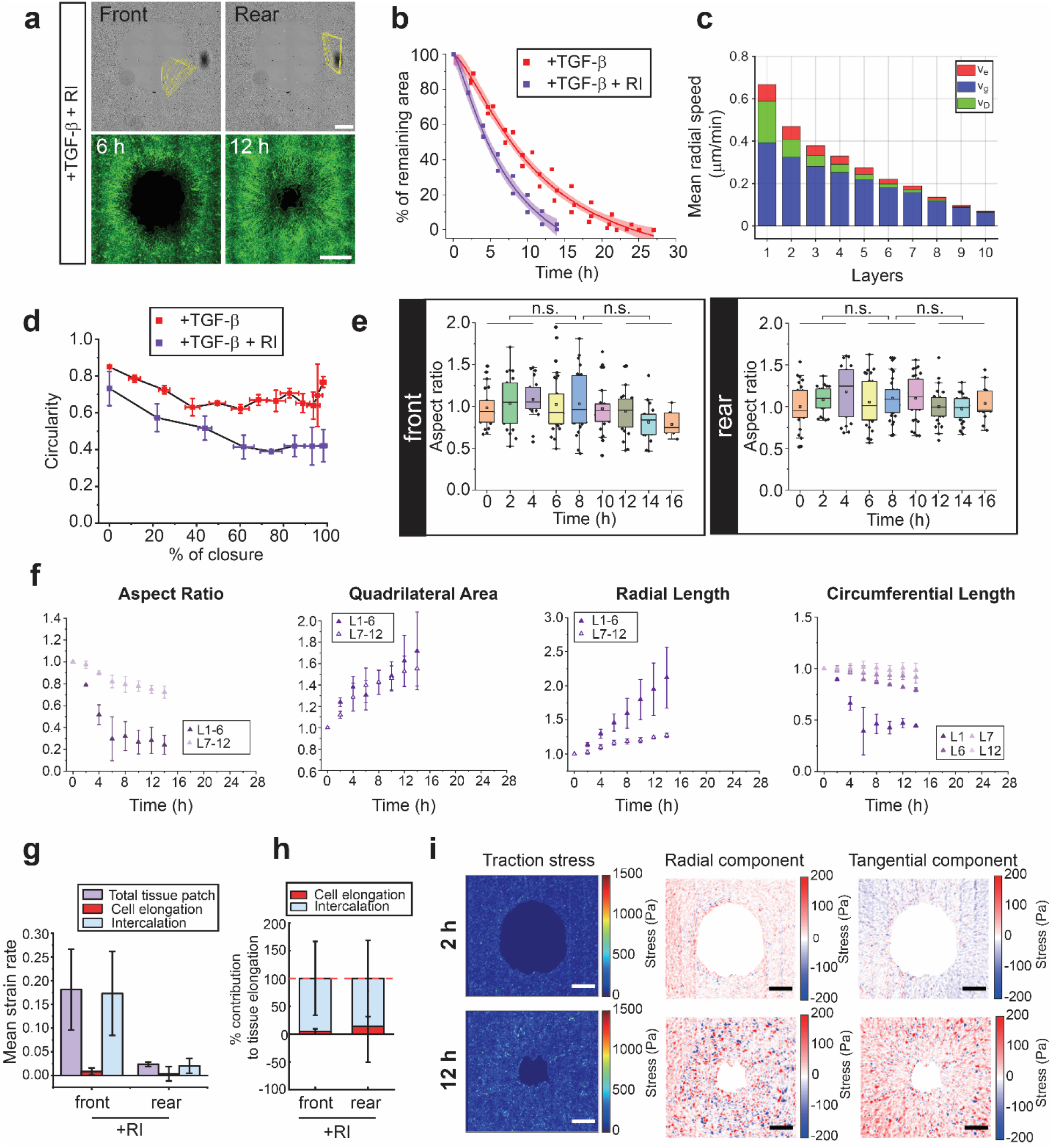
RI increases tissue fluidity and facilitates gap closure. (**a**) Representative brightfield and actin fluorescence images showing the effects of RI treatment on gap closure. Scale bar, 400 µm. (**b**) Plot showing the gap closure dynamics for MEC1 cells treated with or without RI. **(c)** Decomposition of the mean speed of the 10 layers from the best-fit with components due to growth (v_g_), elastic deformation (v_e_), and fluidic rearrangement (v_D_) predicted by the model. (**d**) Plot showing the circularity changes as a function of the remaining percentage of gap area. (**e**) Normalized cell aspect ratio as a function of time. (**f**) Plots showing the tissue patch deformations under RI treatment conditions. (**g-h**) Analysis of mean strain rate of the total tissue patches, and relative contributions from cell elongation and intercalation, respectively. **(i)** Heatmaps showing the magnitude, tangential component, and radial component of traction stress in RI treated MEC1 cells at 2 and 12 h, respectively. Scale bar: 500 μm. Data are represented as mean ± s.e.m. n.s., *P* > 0.05.

## DISCUSSION

In this work we have investigated the cellular and intercellular mechanism for profound gap closure of epicardial cells. We found that during the closure of a circular gap, front 1-2 layers of cells displayed high migration speed and persistence, prominent focal adhesions, and significantly increased cell sizes, regardless of gap sizes. Unlike transient and dynamic nature of leader cell formation mediated by stress-dependent cell polarization found in MDCK cells^13^, leader cells maintained their phenotype throughout the entire process and their migration speed only reduced slightly towards the end. These leader cells were not polarized along radial direction, maintained active proliferation, while having fast and directional motions. The mechanical interactions between leader cells and followers are drastically different from classical epithelial models. While the tensile forces from substrate might drive front layer cell migration, reduced force magnitude in front cells upon TGF-β1 treatment did not attenuate front cell migration speed. In small gap closures, due to small strains and limited time, tensile forces either from substrates or supracellular actin cables are sufficient to deform the monolayer, independent of cell proliferation. For example, inhibiting cell proliferation by mitomycin C treatment did not delay MDCK gap closure within 10 h^51,52^. In contrast, in aphidicolin treated MEC1 cells, the migration speed of the front cells was at similar level as untreated controls but started to steadily drop after 6 h, supporting the essential role of compressive forces from followers in facilitating the leader cell migration. Moreover, for MDCK cells, force generated by leader cells can only influence a relatively small range of about 200 µm (as determined by velocity correlation)^53^. In MEC1 cells in profound gap closure, a larger scale coordinated cell migration and convergent-extension like tissue deformation was observed (7-10 cell layers, about 400-500 µm for 1.7 mm gap), which expanded with decreasing gap curvatures. Regardless of initial gap sizes, we did not observe any purse-string like supracellular actin rings near the gap edge throughout the entire MEC1 gap closure process.

In addition to the unique leader cell behaviors, tissue growth driven mechanism, and tissue-level extension-contraction, we also revealed how EMT contributed to epicardial gap closure at the cell and intercellular level. EMT, particularly in the context to tumorigenesis, is thought to enhance the cell motility, facilitate stress fiber formation, and increase tissue fluidity due to the cadherin switch from stable E-cadherin to transient N-cadherin mediated cell-cell interactions^54,55^. When EMT was induced, we noticed a mildly faster closure in comparison with the control, associated with significant cell elongations in the rear regions and enhanced development of stress fiber radial alignment. Together with tissue-patch deformation analysis, we hypothesized that inducing EMT effectively downregulated tissue fluidity by reducing cell-cell intercalation relative to the observed strain rate (**Fig. 8**).

**Figure 8.**
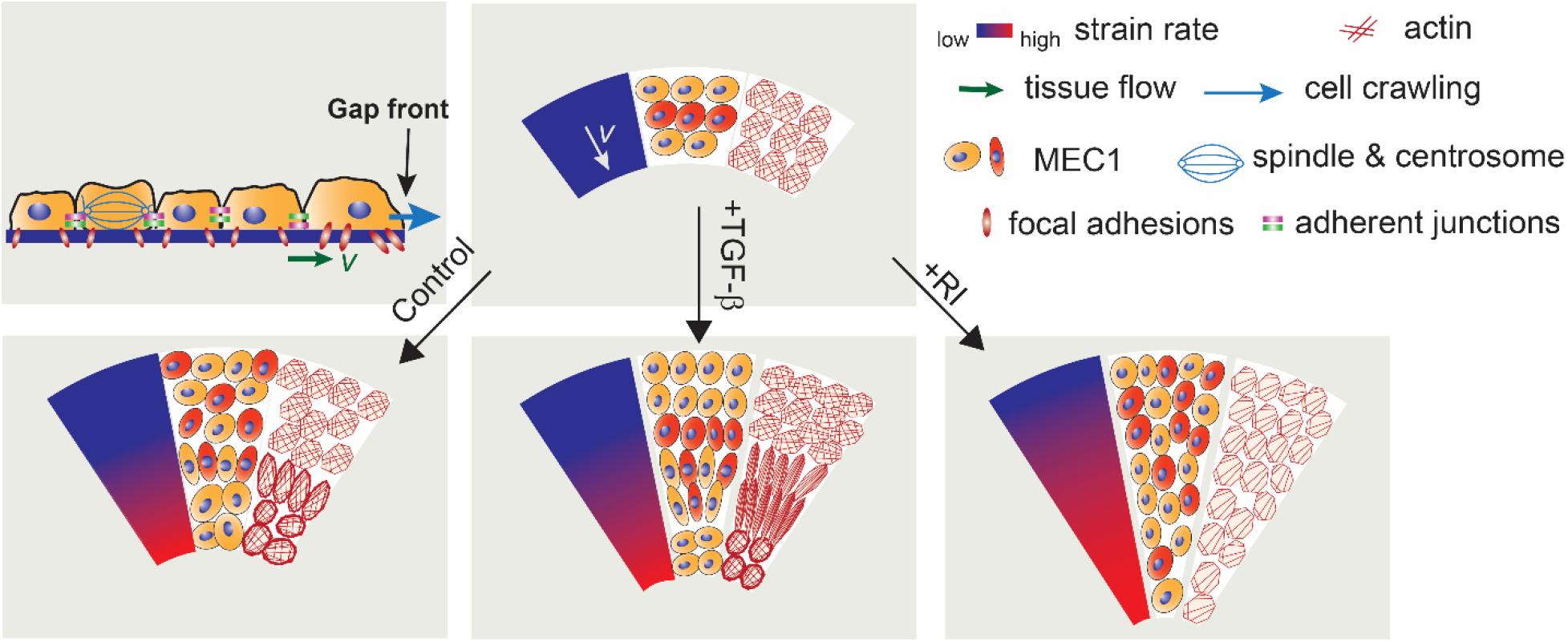
Schematics of the MEC1 gap closure mechanism. In each circular sector, the tissue strain rate gradient (left), cell-cell intercalation (middle), and actin structure and elongation (right) were shown under different conditions. TGF-β1 treatment led to reduced cell-cell intercalation but increased cell elongation. RI treatment increased global cell-cell intercalation and accelerated gap closure. On the top left, cells near the gap front showed increased size and crawled towards the gap center with large focal adhesions. The growth and cell proliferation led to lateral compression in the front of the tissue.

To elucidate such tendency, we developed a Maxwell-type growth viscoelastic model and fit the model parameters to the kinematics data from the two conditions along the partial EMT spectrum. With spatiotemporally uniform parameters, the model simulations were able to capture the key differences between the two conditions qualitatively: the control with more localized inward flow in the front region and less extension-contraction in the rear regions in comparison with the TGF-β1 treated condition. Indeed, the fitted parameters (**Supplementary Table 1**) revealed that the spatiotemporally averaged fluidity tendency in the TGF-β1 treated condition was lower than that in the control. In addition, it revealed that a slightly faster closure was partially attributed to increased cell growth and/or proliferation^56,57^. Further tissue flow and deformation decomposition analysis from the model revealed that the tissue-level extension-contraction was mainly facilitated by elastic deformation in the TGF-β1 treated condition. This result corroborated with more cell elongations when EMT was induced. It is possible that other types of cadherins, such as P-cadherin, play distinct role in stabilizing cell-cell junctions^58^. In addition, our model suggested that the enhanced cell elastic deformation/elongation provided additional directional cues for stress fiber to align radially, which resembled the actomyosin reinforcement along the stretched direction of cells in the Drosophila wing disc^59^.

The role of Rho kinase in collective cell migration is multifaced. In cell systems where cell crawling drives cell migration, RI reduces myosin-mediated contractility and hampers cell migration. It has been shown that RI treatment slowed down gap closure in MDCK cells^11,60^ and fibroblasts^61^. However, it is the opposite case for tension-driven small gap closure due to enhanced cell-cell intercalations near the gap border^17^. For the MEC1 cells, we have also found that RI treatment sped up the closure facilitated by large-scale cell-cell intercalations. Together with the theoretical model, we concluded that RI promoted the growth-driven closure by enhancing global tissue fluidity (**Fig. 8**).

During epithelial morphogenesis, the cell-cell intercalation is promoted by polarized and pulsatile myosin-mediated contractility^12,62^. For more static tissues, however, the cell-cell intercalation is promoted by weakened contractility^63^. With the partial EMT phenotype, the MEC1 cell monolayer presents strikingly different medial and intercellular actin organization. Thus, the previous cell-level understanding of cell-cell intercalation due to intercellular tension^23,64,65^ cannot be directly applied here. One of the future works is to understand the cell-cell intercalation at the cell level, by developing a more general vertex model integrated with the continuum mechanics. Such a model would account for tissue growth and the actin fiber organization associated with the partial EMT phenotype, with both represented in a manner consistent with cell growth and division and cell-specific actin remodeling. In particular, we propose that the change of cell sizes in zones 1 and 2 reflect both the cell-cycle progression and response to the growing planar compressive stress predicted by the continuum model^48^. Such hypothesis would be examined within the integrated modeling approach.

Besides revealing that the inward migration is driven by compressive forces within the growing monolayer, the traction maps also reveal striking differences among the control, the TGF-β1 – treated, and the TGF-β1/RI–treated conditions. For the TGF-β1-treated case, the cell–substrate interaction is the strongest spatiotemporally and persists even after gap closure. For the TGF-β1/RI–treated condition, the interaction is minimal. Thus, the reduced obstructive force from the substrate may also contribute to the faster closure compared with the other two cases. Due to the complex spatiotemporal patterning of the traction forces (with directions changing between the front and rear regions), we chose to construct a minimal model that did not explicitly incorporate traction patterning. Future work will refine the study to regional resolutions and elucidate the differences in mechanical interactions between the front and rear regions.

In summary, in this work we revealed a novel gap closure mechanism by epicardial cells that is primarily driven by growth-mediated compressive stresses and facilitated by large-scale tissue extension contraction. TGF-β1-induced partial EMT, caused accelerated gap closure, relatively less cell-cell intercalation, and tissue-wide actin re-organization associated with cell elongation. Our results reveal new modes of tissue fluidity regulation and inspire novel intervention strategies targeting proliferation and tissue fluidity, with potential applications ranging from cardiac tissue regeneration to a broader class of tissue migration processes involving partial EMT.

## METHODS

### Cell culture

MEC1 cells were cultured in the Dulbecco’s modified Eagle’s medium (DMEM; Gibco, 11960051), with 10% fetal bovine serum (FBS; Sigma or Cytiva, SH30071.03), 1% GlutaMax (Gibco, 35050061), 1% Sodium Pyruvate (Gibco, 11360070), 1% MEM NEAA (Non-Essential Amino Acid; Gibco, 11140050), and 1% penicillin/streptomycin (Gibco, 15140122); MDCK cell were cultured in the DMEM with 10% FBS, 1% GlutaMax, 1% Sodium Pyruvate, and 1% penicillin/streptomycin. All cells were cultured in a cell culture incubator at 37°C with 5% CO_2_. The MEC1 cell line was obtained from Dr. Jian Xu’s lab. MDCK cells were purchased from ATCC.

### Gap generation

PDMS stencils were generated using PDMS soft lithography using 3D printed molds (ProFluidics 285D). Dishes and PDMS stencils were treated with Ultraviolet Ozone Cleaner (Jelight) for 7 minutes. After treatment, dishes were coated with Fibronectin (10 µg/mL), and PDMS stencils were coated with Pluronic® F-127 (4% v/w; Sigma, P2443-250G) for 1 hour. Dishes and stencils were rinsed with DI water and blown dry with an electronic duster. The stencil was put on the coverslip bottom of the fibronectin-coated dish and the cell suspension solution was injected under the stencil with micropipettes. After 40 mins of incubation, 2 mL of culture media were added into the dish. Cells were cultured with the stencil in the incubator overnight before the stencil was removed.

### Immunocytochemistry

Cells were fixed with 4% paraformaldehyde and permeabilized with 0.1% Triton X-100. Samples were blocked with 10% donkey serum for 1 hour at room temperature and incubated with the following primary antibodies: anti-YAP (Abcam, sc-101199), anti-Phospho-Histone H3 (Cell Signaling Technology, 53348T), and anti–Golgin-97 (Invitrogen, A21270). For immunolabeling, donkey-anti rabbit Alexa Fluor 555 and donkey-anti mouse Alexa Fluor 647 were used. For actin filaments visualization, Alexa Fluor 488 conjugated phalloidin (Invitrogen, A12379) was used. Samples were counterstained with 4,6-diamidino-2-phenylindole (DAPI; Invitrogen). Images were acquired using inverted epifluorescence microscope (Leica DMi8; Leica Microsystems) equipped with a monochrome charge-coupled device (CCD) camera.

### Live cell imaging

Time-lapse video-microscopy was performed by Cytation 3 Cell Imaging Multi-Mode Reader (BioTek). Cells were cultured in 35 mm glass bottom dishes at 37 °C and with 5% CO_2_. Cells were imaged for 30 hours after stencil removal. Images were obtained every 20 min for the indicated time courses.

### Traction force microscopy

Coverslips were treated with 1% APTMS (Sigma Aldrich 21788) for 15 minutes, rinsed and incubated with 0.5% Glutaraldehyde solution overnight. Then, the coverslip was mounted in a modified MatTek 35 mm dish with a 20 mm hole. 200 nm red fluorescent polystyrene microspheres (Life Technologies, F8810) were dispersed in pure ethanol at a ratio of 1:200 and spun coated on plasma treated coverslips twice at 5,000 rpm for 20 seconds. Polyacrylamide (PAA) solution (12% acrylamide and 0.14% bisacrylamide in 50 mM HEPES) were added with 0.1% ammonium persulfate (Amresco, Solon, OH, USA, Cat. No. 7727540) and 0.3% N,N,N′,N′-tetramethylethylenediamine (Amresco, Cat. No. 110189) and dropped of the solution on the center of the coverslip with beads (48 μL/coverslip). The prepared dish was gently placed onto the droplet to allow the polyacrylamide solution to spread uniformly between the coverslips. The solution was polymerized for 15 min at room temperature to form a 20-kPa polyacrylamide gel disk (20 mm diameter, 150 μm thickness) with fluorescent beads embedded ∼5 μm below the gel surface. Collagen type I was cross-linked to the gel surface using sulfo-SANPAH (Proteochem, 102568-43-4). Gels were submerged in 1 mg·mL⁻¹ sulfo-SANPAH and exposed to UV light (8 W) for 15 min, washed with HEPES buffer, and incubated with 0.1 mg·mL⁻¹ rat-tail collagen type I (Corning, 354249) for 12 h at 4 °C. Collagen-coated gels were washed with PBS buffer, equilibrated in culture medium for 30 min at 37 °C, and used for cell gap generation.

Cells and fluorescent beads were imaged at 2, 12, and 24 h after stencil removal. At each time point, a mosaic of 9 × 11 images with 15% overlap was acquired, distortion-corrected, and stitched to generate a high-resolution image of the entire gap region and the underlying fluorescent beads. After removal of cells with 0.05% trypsin, an additional set of bead images was acquired and stitched to generate a reference image of the undeformed gel. Images were processed in MATLAB using single-particle tracking to extract bead centroids. Gel surface deformation at each time point was quantified from bead displacements, calculated as the difference between bead centroids and their positions in the reference image. Cell traction forces were computed from the gel surface deformation using finite element analysis (ANSYS)^66^.

### Actin orientation structure parameter

To describe the orientation of the actin fiber at each location along the radius, we considered a structure tensor: 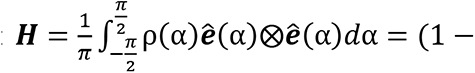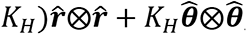, where 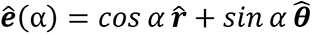 , the unit vector represented the fiber orientation with α the angle between the actin fiber and radial directions, *ρ*(*α*) denoted the angle density distribution 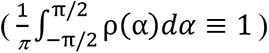 of the actin fiber orientation, and the structural parameter 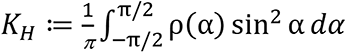. For the experimental data (Fig.1h and 2h), *K_H_* was computed after extracting the distribution ρ(α) from the software developed in our previous work^39^. For the simulation (Fig.6h), we assumed the initial density *ρ*^0^ ≡ 1 was uniform from the fact that 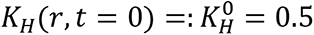 (**Supplementary Fig. 5**), and computed 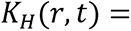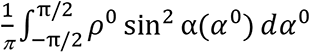, where the mapping 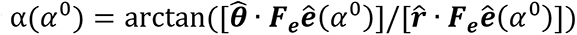 by assuming that the elastic deformation ***F_e_*** changes the local fiber orientation as affine transformation.

### Particle Image Velocimetry

The velocity field of the gap closure at different times were characterized using the PIVlab, an open-source particle image velocimetry (PIV) MATLAB tool (https://www.mathworks.com/matlabcentral/fileexchange/27659-pivlab-particle-image-velocimetry-piv-tool-with-gui/). Bright-field image series showing the gap closure process with 20 minutes time intervals were pre-processed to increase the contrast. The displacements from position to position, and hence the velocity field at that time were computed using the PIVlab. The strain rate field 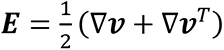 in the tissue was computed using central difference while filtering out data points next to the gap border. The growth rate *tr**E*** and the spectral radius of 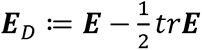 fields were computed algebraically at each point. 2D regions were divided into quasi-concentric layers using signed distance from the gap border, then we averaged each 2D-field variable on each layer and plotted it as a function of the radial position.

### Pharmacological treatment

For TGF-β1 treatment, a stock solution of TGF-β1 (5µg/mL, Proteintech) was prepared by dissolving TGF-β1 in 4 mM HCl water solution with 0.1% BSA (50 mg/mL). The stock solution was diluted in culture media to a working concentration of 2 ng/mL. For Aphidicolin treatment, a stock solution (1 mg/mL) was prepared using DMSO as solvent. The stock solution was then diluted to a working concentration of 1 μg/mL in culture media. For Y27632 (RI) treatment, a stock solution of Y27632 (10 mM) was prepared in water and diluted to a working concentration of 10 μM in culture media.

### Tissue deformation quantification

To quantify tissue deformation dynamics, we defined a geometric reference by randomly selecting two cells from each of layers 1 and 5, or layers 6 and 10. The four cells in layers 1 and 5 represented the front region, while the cells in layers 6 and 10 represented the rear region. A quadrilateral reference shape was formed with these four cells as vertices, and this quadrilateral was used to represent the tissue region. After generating the quadrilateral, the line formed by the two cells in layer 1 was recorded as the circumferential length L1, the line in layer 5 was recorded as the circumferential length L5, the average value of the two sides was recorded as the radial length; and the ratio of the smaller circumferential length (i.e., L1 and L6) to the radial length was defined as the aspect ratio. The migration trajectories of the vertex positions enabled continuous reconstruction of the quadrilateral geometry and tracking the shape throughout the closure process. All the data were normalized to 0 h values to mitigate the variation in initial quadrilateral sizes.

### Cell proliferation rate calculation

The cell proliferation rate, *r*, is calculated as: *r* = 1/*t_m_* · ln(1 + 2*R*)/(1 + *R*); where *R* is the fraction of cells undergoing mitosis; *t_m_* is the time required for mitosis. In our experiments, *R* was determined by the percentage of the pHH3 positive cells and *t_m_* was obtained from bright-field video microscopy. Briefly, bright-field images were captured at 20-minute intervals. Cells approaching mitosis were identified and tracked based on morphological features and *t_m_* was calculated by averaging the estimated time for cell mitosis at different locations in three samples.

### Statistics

Statistical analysis was performed using GraphPad Prism. Data were tested for normality. For statistical comparations between two data sets, *P*-values were calculated using the student *t*-test function or the Mann-Whitney test when appropriate. For statistical comparations between three or more data sets, *P*-values were calculated using the one-way ANOVA with Tukey post-hoc analysis or the Kruskal-Wallis test when appropriate.

### Analysis of cell and tissue patch aspect ratio rate

The aspect ratio for both cells *λ_cell_* and tissue patches λ are evaluated) are evaluated with the circumferential dimension over the radial dimension. Both *λ_cell_* and λ are normalized by the initial condition. The mean rate of change of the aspect ratio of tissue patches (cells) is determined by 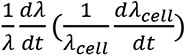. The difference 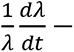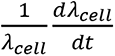 is the rate contribution from the cell intercalation.

### Simulation of cell layer migration and tissue-patch deformation

To decompose the tissue flow and deformation, for any given velocity 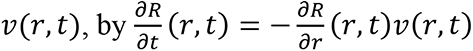 with initial value *R*(*r*, 0) = *R*_0_, we could find the reference mapping *R*(*r*, *t*) at time *t*. By the inverse of reference mapping, we could then find the migration of each layer by interpolation {*r_i_*(*t*): *R*(*r_i_*(*t*), *t*) = *R_i_*}, where *R_i_* is the initial radius of *i*-th layer. Next, the velocity of each layer was given by *v*(*r_i_*(*t*), *t*). Similarly, we could find the stresses *σ*(*r_i_*(*t*), *t*) and elastic deformation *F_e_* at each layer. We decomposed the mean speed of the first ten layers of cells (*v*) from the best-fit with components due to growth (*v_g_*), elastic deformation (*v_e_*), and fluidic rearrangement (*v_D_*), and *v* = *v_g_* + *v_e_* + *v* . We started from the decomposition for strain rates: 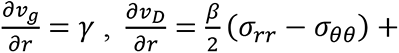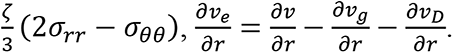. For *i*-th layer, we obtained *v* (*t*) by numerical integration, 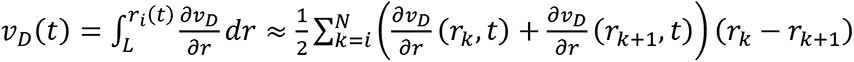 , where *r _N+1_*= *L* is the fixed boundary where *v* = 0 . Similarly, we could obtain *v_g_*(*t*) = *γ* ⋅ (*r_i_*(*t*) − *L*) , and hence *v_e_*(*t*) = *v* − *v_g_* − *v_D_*.

To calculate the area and aspect ratio changes as a function of time, the total aspect ratio of *i*-th layer is defined as *λ* = *λ_θ_*/*λ_r_*, where 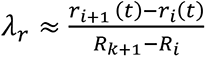 are the stretch in the radial and circumferential directions, respectively. The elastic aspect ratio is given by *λ_e_* = *f_er_*(*r_i_*(*t*))/*f_eθ_*(*r_i_*(*t*)) , and then the fluidic aspect ratio is *λ_D_* = *λ*/*λ_e_* given that *λ_g_* = exp(*γt*)/ exp(*γt*) = 1, where exp(*γt*) is the growth stretch in both radial and circumferential directions with in-plane isotropic growth rate *γ*.

The area change of *i*-th layer is *J* = *λ_r_* ⋅ *λ_θ_*. Similarly, the elastic area change is *J_e_* = *f_er_*(*r_i_*(*t*)) ⋅ *f_eθ_*(*r_i_*(*t*)) and the fluidic area change is *J_D_* = *J*/(*J_g_J_e_*), where *J_g_* = exp(2*γt*).

## AUTHOR CONTRIBUTIONS

Y.S. and M.W. conceived and planned the project and wrote the manuscript with support from all of the authors. H.J. performed experiments and collected and analyzed data. C. W. performed simulations and mathematical modeling. P.W. and Q.W. performed traction force microscopy analysis. J. J. performed focal adhesion staining. N.O. and Y.G. performed PIV analysis. C. W., F.C., and J.X. contributed to data analysis. Funding was acquired by Y. S., M. W. and Q. W.

## ACKNOWLEDGMENT

This work is supported in part by the National Institute of Health (R35GM155279 and R01DK129990 to Y.S. and R01GM157590 to M. W. and Q. W.). M. W. additionally acknowledges partial funding support by the National Science Foundation – Division of Mathematical Sciences (NSF-DMS) under grants DMS-2012330 and DMS-2144372. We acknowledge the ADDFab and Light Microscopy Facilities at UMass Amherst for 3D printing and microscopy.

## REFERENCES

1 Stossel, T. P. On the crawling of animal cells. Science 260, 1086–1094, doi:10.1126/science.8493552 (1993).

2 Martin, P. Wound Healing--Aiming for Perfect Skin Regeneration. Science 276, 75–81, doi:10.1126/science.276.5309.75 (1997).

3 Martin, P. & Lewis, J. Actin cables and epidermal movement in embryonic wound healing. Nature 360, 179–183, doi:10.1038/360179a0 (1992).

4 Bement, W. M., Forscher, P. & Mooseker, M. S. A novel cytoskeletal structure involved in purse string wound closure and cell polarity maintenance. J Cell Biol 121, 565–578, doi:10.1083/jcb.121.3.565 (1993).

5 Gipson, I. K. & Danjo, Y. Actin ‘purse string’ filaments are anchored by E-cadherin-mediated adherens junctions at the leading edge of the epithelial wound, providing coordinated cell movement. J Cell Sci 111 (1998).

6 Williams-Masson, E. M., Malik, A. N. & Hardin, J. An actin-mediated two-step mechanism is required for ventral enclosure of the C. elegans hypodermis. Development 124, 2889–2901 (1997).

7 Young, P. E., Richman, A. M., Ketchum, A. S. & Kiehart, D. P. Morphogenesis in Drosophila requires nonmuscle myosin heavy chain function. Genes Dev 7, 29–41, doi:10.1101/gad.7.1.29 (1993).

8 Anon, E. et al. Cell crawling mediates collective cell migration to close undamaged epithelial gaps. Proceedings of the National Academy of Sciences 109, 10891–10896, doi:10.1073/pnas.1117814109 (2012).

9 Klarlund, J. K. Dual modes of motility at the leading edge of migrating epithelial cell sheets. Proceedings of the National Academy of Sciences 109, 15799–15804, doi:10.1073/pnas.1210992109 (2012).

10 Wei, Q. et al. Actin-ring segment switching drives nonadhesive gap closure. Proceedings of the National Academy of Sciences, 202010960, doi:10.1073/pnas.2010960117 (2020).

11 Fenteany, G., Janmey, P. A. & Stossel, T. P. Signaling pathways and cell mechanics involved in wound closure by epithelial cell sheets. Curr Biol 10, 831–838, doi:10.1016/s0960-9822(00)00579-0 (2000).

12 Brugues, A. et al. Forces driving epithelial wound healing. Nat Phys 10, 684–691, doi:10.1038/Nphys3040 (2014).

13 Vishwakarma, M. et al. Mechanical interactions among followers determine the emergence of leaders in migrating epithelial cell collectives. Nature Communications 9, 3469, doi:10.1038/s41467-018-05927-6 (2018).

14 Hino, N. et al. A feedback loop between lamellipodial extension and HGF-ERK signaling specifies leader cells during collective cell migration. Developmental Cell 57, 2290–2304.e2297, 10.1016/j.devcel.2022.09.003 (2022).

15 Kozyrska, K. et al. p53 directs leader cell behavior, migration, and clearance during epithelial repair. Science 375, eabl8876, doi:10.1126/science.abl8876 (2022).

16 Riahi, R. et al. Notch1–Dll4 signalling and mechanical force regulate leader cell formation during collective cell migration. Nature Communications 6, 6556, doi:10.1038/ncomms7556 (2015).

17 Tetley, R. J. et al. Tissue Fluidity Promotes Epithelial Wound Healing. Nat Phys 15, 1195–1203, doi:10.1038/s41567-019-0618-1 (2019).

18 Jain, A. et al. Regionalized tissue fluidization is required for epithelial gap closure during insect gastrulation. Nature Communications 11, 5604, doi:10.1038/s41467-020-19356-x (2020).

19 Shindo, A. Models of convergent extension during morphogenesis. Wiley Interdiscip Rev Dev Biol 7, doi:10.1002/wdev.293 (2018).

20 Huebner, R. J. & Wallingford, J. B. Coming to Consensus: A Unifying Model Emerges for Convergent Extension. Dev Cell 46, 389–396, doi:10.1016/j.devcel.2018.08.003 (2018).

21 Wang, X. et al. Anisotropy links cell shapes to tissue flow during convergent extension. Proc Natl Acad Sci U S A 117, 13541–13551, doi:10.1073/pnas.1916418117 (2020).

22 Claussen, N. H., Brauns, F. & Shraiman, B. I. A geometric-tension-dynamics model of epithelial convergent extension. Proc Natl Acad Sci U S A 121, e2321928121, doi:10.1073/pnas.2321928121 (2024).

23 Brauns, F., Claussen, N. H., Lefebvre, M. F., Wieschaus, E. F. & Shraiman, B. I. The geometric basis of epithelial convergent extension. Elife 13, doi:10.7554/eLife.95521 (2024).

24 (!!! INVALID CITATION !!! 17,18).

25 Jackson-Weaver, O. et al. PRMT1-p53 Pathway Controls Epicardial EMT and Invasion. Cell Rep 31, 107739, doi:10.1016/j.celrep.2020.107739 (2020).

26 Wang, J. H., Cao, J. L., Dickson, A. L. & Poss, K. D. Epicardial regeneration is guided by cardiac outflow tract and Hedgehog signalling. Nature 522, 226-+, doi:10.1038/nature14325 (2015).

27 van Wijk, B., Gunst, Q. D., Moorman, A. F. & van den Hoff, M. J. Cardiac regeneration from activated epicardium. PLoS One 7, e44692, doi:10.1371/journal.pone.0044692 (2012).

28 Krainock, M. et al. Epicardial Epithelial-to-Mesenchymal Transition in Heart Development and Disease. J Clin Med 5, doi:10.3390/jcm5020027 (2016).

29 Quijada, P., Trembley, M. A. & Small, E. M. The Role of the Epicardium During Heart Development and Repair. Circ Res 126, 377–394, doi:10.1161/CIRCRESAHA.119.315857 (2020).

30 Campbell, K. et al. Collective cell migration and metastases induced by an epithelial-to-mesenchymal transition in Drosophila intestinal tumors. Nat Commun 10, 2311, doi:10.1038/s41467-019-10269-y (2019).

31 Aiello, N. M. et al. EMT Subtype Influences Epithelial Plasticity and Mode of Cell Migration. Dev Cell 45, 681–695.e684, doi:10.1016/j.devcel.2018.05.027 (2018).

32 von Gise, A. & Pu, W. T. Endocardial and Epicardial Epithelial to Mesenchymal Transitions in Heart Development and Disease. Circ.Res. 110, 1628–1645, doi:10.1161/Circresaha.111.259960 (2012).

33 Rodriguez, E. K., Hoger, A. & McCulloch, A. D. Stress-dependent finite growth in soft elastic tissues. J Biomech 27, 455–467 (1994).

34 Dicko, M. et al. Geometry can provide long-range mechanical guidance for embryogenesis. PLOS Computational Biology 13, e1005443, doi:10.1371/journal.pcbi.1005443 (2017).

35 Streichan, S. J., Lefebvre, M. F., Noll, N., Wieschaus, E. F. & Shraiman, B. I. Global morphogenetic flow is accurately predicted by the spatial distribution of myosin motors. eLife 7, e27454, doi:10.7554/eLife.27454 (2018).

36 Ioratim-Uba, A., Liverpool, T. B. & Henkes, S. Mechanochemical Active Feedback Generates Convergence Extension in Epithelial Tissue. Physical Review Letters 131, 238301, doi:10.1103/PhysRevLett.131.238301 (2023).

37 Saadaoui, M., Rocancourt, D., Roussel, J., Corson, F. & Gros, J. A tensile ring drives tissue flows to shape the gastrulating amniote embryo. Science 367, 453–458, doi:10.1126/science.aaw1965 (2020).

38 Wu, M. & Ben Amar, M. Growth and remodelling for profound circular wounds in skin. Biomech Model Mechanobiol 14, 357–370, doi:10.1007/s10237-014-0609-1 (2015).

39 Xie, T. et al. Condensation tendency and planar isotropic actin gradient induce radial alignment in confined monolayers. eLife 10, e60381, doi:10.7554/eLife.60381 (2021).

40 Cao, J. L. et al. Tension Creates an Endoreplication Wavefront that Leads Regeneration of Epicardial Tissue. Developmental Cell 42, 600-+, doi:10.1016/j.devcel.2017.08.024 (2017).

41 Dupont, S. et al. Role of YAP/TAZ in mechanotransduction. Nature 474, 179–183, doi:10.1038/nature10137 nature10137 [pii] (2011).

42 Khalil, A. A. et al. A YAP-centered mechanotransduction loop drives collective breast cancer cell invasion. Nature Communications 15, 4866, doi:10.1038/s41467-024-49230-z (2024).

43 Kim, D., Olson, J. M. & Cooper, J. A. N-cadherin dynamically regulates pediatric glioma cell migration in complex environments. Journal of Cell Biology 223, doi:10.1083/jcb.202401057 (2024).

44 Chung, K. T., Nilson, E. H., Case, M. J., Marr, A. G. & Hungate, R. E. Estimation of growth rate from the mitotic index. Appl Microbiol 25, 778–780, doi:10.1128/am.25.5.778-780.1973 (1973).

45 Pedrali-Noy, G. et al. Synchronization of HeLa cell cultures by inhibition of DNA polymerase alpha with aphidicolin. Nucleic Acids Res 8, 377–387, doi:10.1093/nar/8.2.377 (1980).

46 Wallingford, J. B., Fraser, S. E. & Harland, R. M. Convergent extension: the molecular control of polarized cell movement during embryonic development. Dev Cell 2, 695–706 (2002).

47 Kupfer, A., Louvard, D. & Singer, S. J. Polarization of the Golgi apparatus and the microtubule-organizing center in cultured fibroblasts at the edge of an experimental wound. Proc Natl Acad Sci U S A 79, 2603–2607, doi:10.1073/pnas.79.8.2603 (1982).

48. Wei, C., et al. Continuum modeling of fluidic and elastic flow during growth-driven wound closure in partial-EMT cell monolayers. arXiv, https://arxiv.org/abs/2607.05820 (2026).

49 Ranft, J. et al. Fluidization of tissues by cell division and apoptosis. Proceedings of the National Academy of Sciences 107, 20863–20868, doi:10.1073/pnas.1011086107 (2010).

50 Melchionna, R., Trono, P., Tocci, A. & Nisticò, P. Actin Cytoskeleton and Regulation of TGFβ Signaling: Exploring Their Links. Biomolecules 11, doi:10.3390/biom11020336 (2021).

51 Farooqui, R. & Fenteany, G. Multiple rows of cells behind an epithelial wound edge extend cryptic lamellipodia to collectively drive cell-sheet movement. J Cell Sci 118, 51–63, doi:10.1242/jcs.01577 (2005).

52 Poujade, M. et al. Collective migration of an epithelial monolayer in response to a model wound. Proc Natl Acad Sci U S A 104, doi:10.1073/pnas.0705062104 (2007).

53 Petitjean, L. et al. Velocity fields in a collectively migrating epithelium. Biophys J 98, 1790–1800, doi:10.1016/j.bpj.2010.01.030 (2010).

54 Revenu, C. & Gilmour, D. EMT 2.0: shaping epithelia through collective migration. Current Opinion in Genetics & Development 19, 338–342, 10.1016/j.gde.2009.04.007 (2009).

55 Kuriyama, S. et al. In vivo collective cell migration requires an LPAR2-dependent increase in tissue fluidity. J Cell Biol 206, 113–127, doi:10.1083/jcb.201402093 (2014).

56 Lamouille, S. & Derynck, R. Cell size and invasion in TGF-beta-induced epithelial to mesenchymal transition is regulated by activation of the mTOR pathway. J Cell Biol 178, 437–451, doi:10.1083/jcb.200611146 (2007).

57 Hosseini, K., Trus, P., Frenzel, A., Werner, C. & Fischer-Friedrich, E. Skin epithelial cells change their mechanics and proliferation upon snail-mediated EMT signalling. Soft Matter 18, 2585–2596, doi:10.1039/D2SM00159D (2022).

58 Dang, Q. M., Smart, N., Redpath, A. N. & Vieira, J. M. Organ-specific and conserved regulatory logic orchestrates gene expression in the embryonic mesothelium. bioRxiv, 2025.2008.2006.668886, doi:10.1101/2025.08.06.668886 (2025).

59 LeGoff, L., Rouault, H. & Lecuit, T. A global pattern of mechanical stress polarizes cell divisions and cell shape in the growing Drosophila wing disc. Development 140, 4051–4059, doi:10.1242/dev.090878 (2013).

60 Isozaki, Y. et al. The Rho-guanine nucleotide exchange factor Solo decelerates collective cell migration by modulating the Rho-ROCK pathway and keratin networks. Mol Biol Cell 31, 741–752, doi:10.1091/mbc.E19-07-0357 (2020).

61 Sakar, M. S. et al. Cellular forces and matrix assembly coordinate fibrous tissue repair. Nature Communications 7, doi:ARTN 11036 10.1038/ncomms11036 (2016).

62 Collinet, C., Rauzi, M., Lenne, P. F. & Lecuit, T. Local and tissue-scale forces drive oriented junction growth during tissue extension. Nat Cell Biol 17, 1247–1258, doi:10.1038/ncb3226 (2015).

63 Curran, S. et al. Myosin II Controls Junction Fluctuations to Guide Epithelial Tissue Ordering. Dev Cell 43, 480–492 e486, doi:10.1016/j.devcel.2017.09.018 (2017).

64 Bi, D., Lopez, J. H., Schwarz, J. M. & Manning, M. L. Energy barriers and cell migration in densely packed tissues. Soft Matter 10, 1885–1890, doi:10.1039/c3sm52893f (2014).

65 Bi, D., Lopez, J. H., Schwarz, J. M. & Manning, M. L. A density-independent rigidity transition in biological tissues. Nat Phys 11, 1074–1079, doi:10.1038/nphys3471 (2015).

66 Ho Thanh, M.-T., Grella, A., Kole, D., Ambady, S. & Wen, Q. Vimentin intermediate filaments modulate cell traction force but not cell sensitivity to substrate stiffness. Cytoskeleton 78, 293–302, 10.1002/cm.21675 (2021).

