## Supplementary figure and table for "Partial epithelial-to-mesenchymal transition mediates profound gap closure through growth and fluidization"

**Supplementary Figure 1**

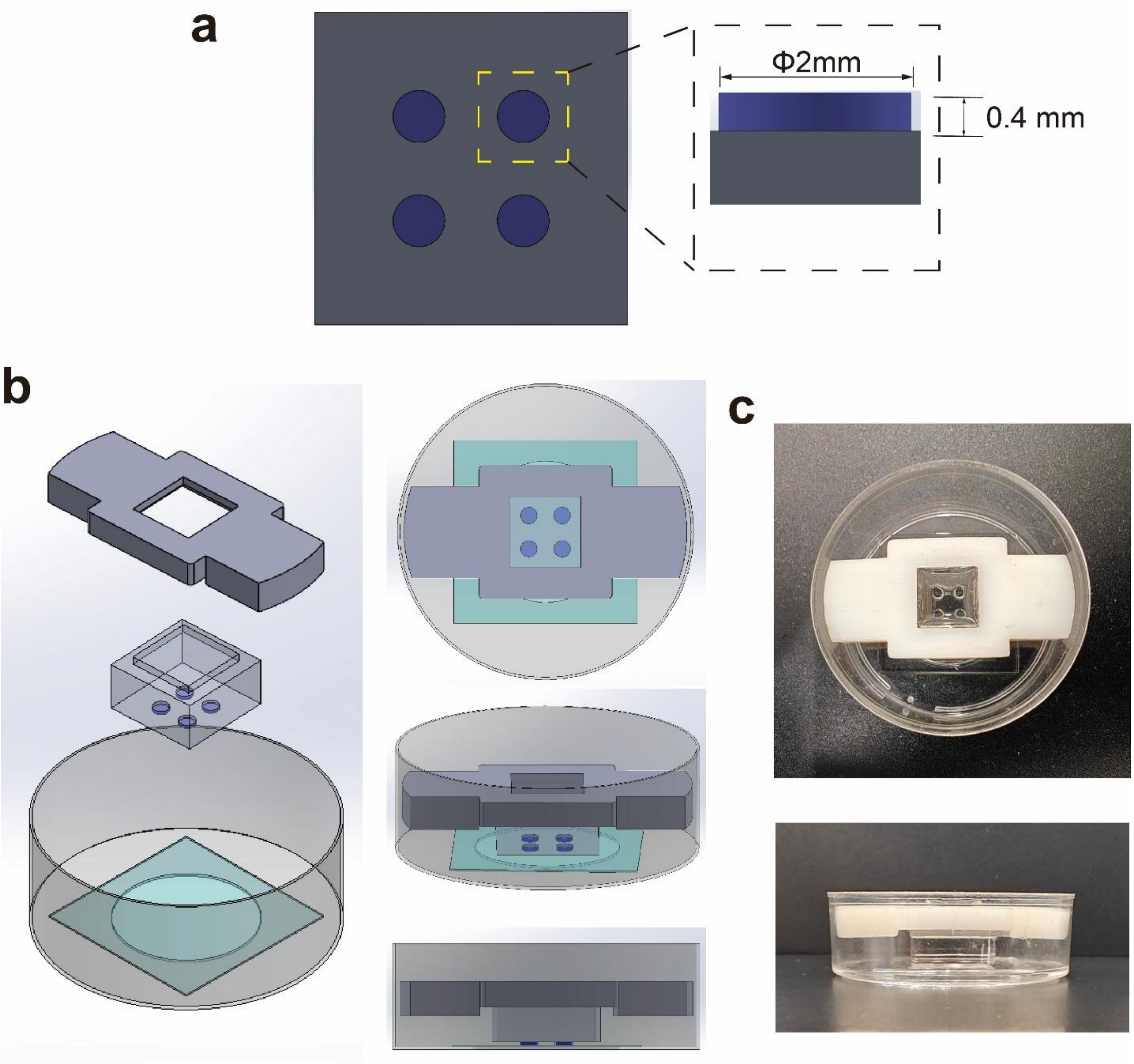


**Supplementary Fig. 1.** Microdevices for generating controlled gaps. (**a**) Design of the stencils. (**b**) Schematics showing the assembly of the stencil and mounting device for generating large circular gaps on coverslips. The stencils and mounting device were 3D printed (**c**) Photographs showing top and side views of the experimental setup.

**Supplementary Figure 2**

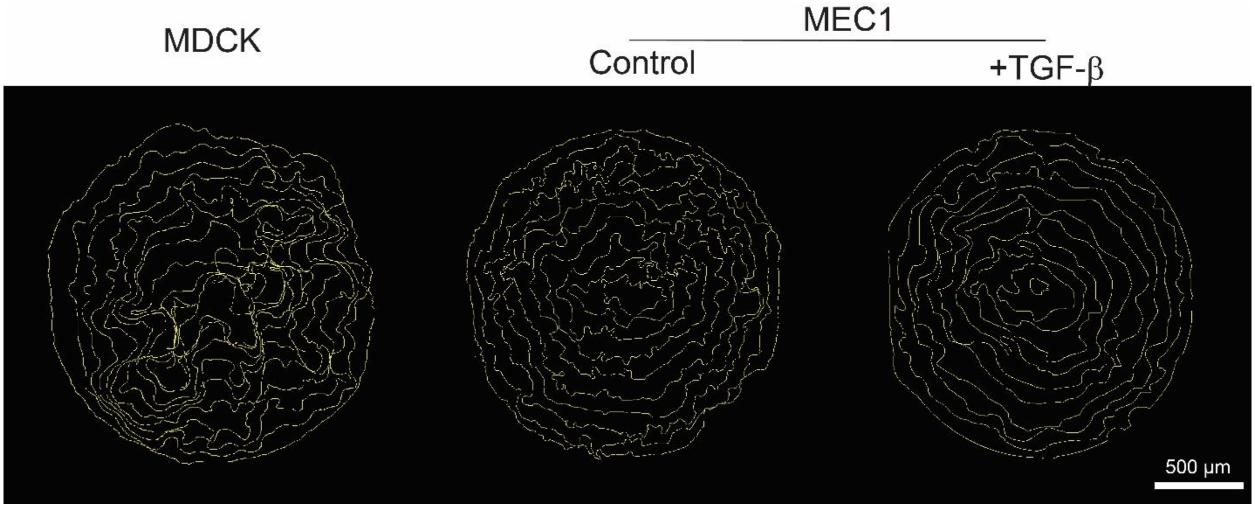


**Supplementary Fig. 2.** Traces of gap fronts. Timelapse videomicroscopy was performed and the front of the gap was traced every 2 hours. Scale bar, 500 µm.

**Supplementary Figure 3**

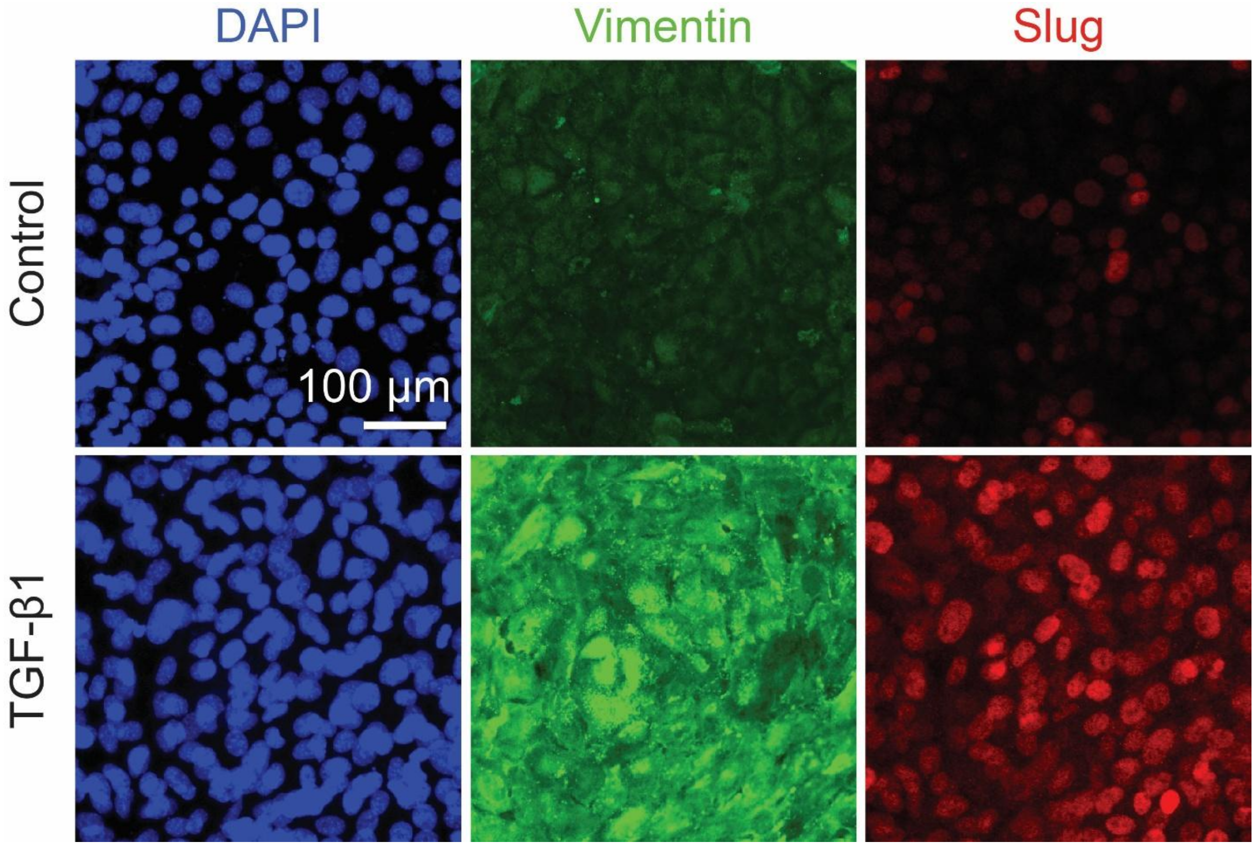


**Supplementary Fig. 3.** TGF-β1 treatment induced EMT in MEC1 cells. Fluorescence images showing the activation of EMT markers, including vimentin and slug, in MEC1 cells after TGF-β1 treatment for 24 hours. Scale bar, 100 μm.

**Supplementary Figure 4**

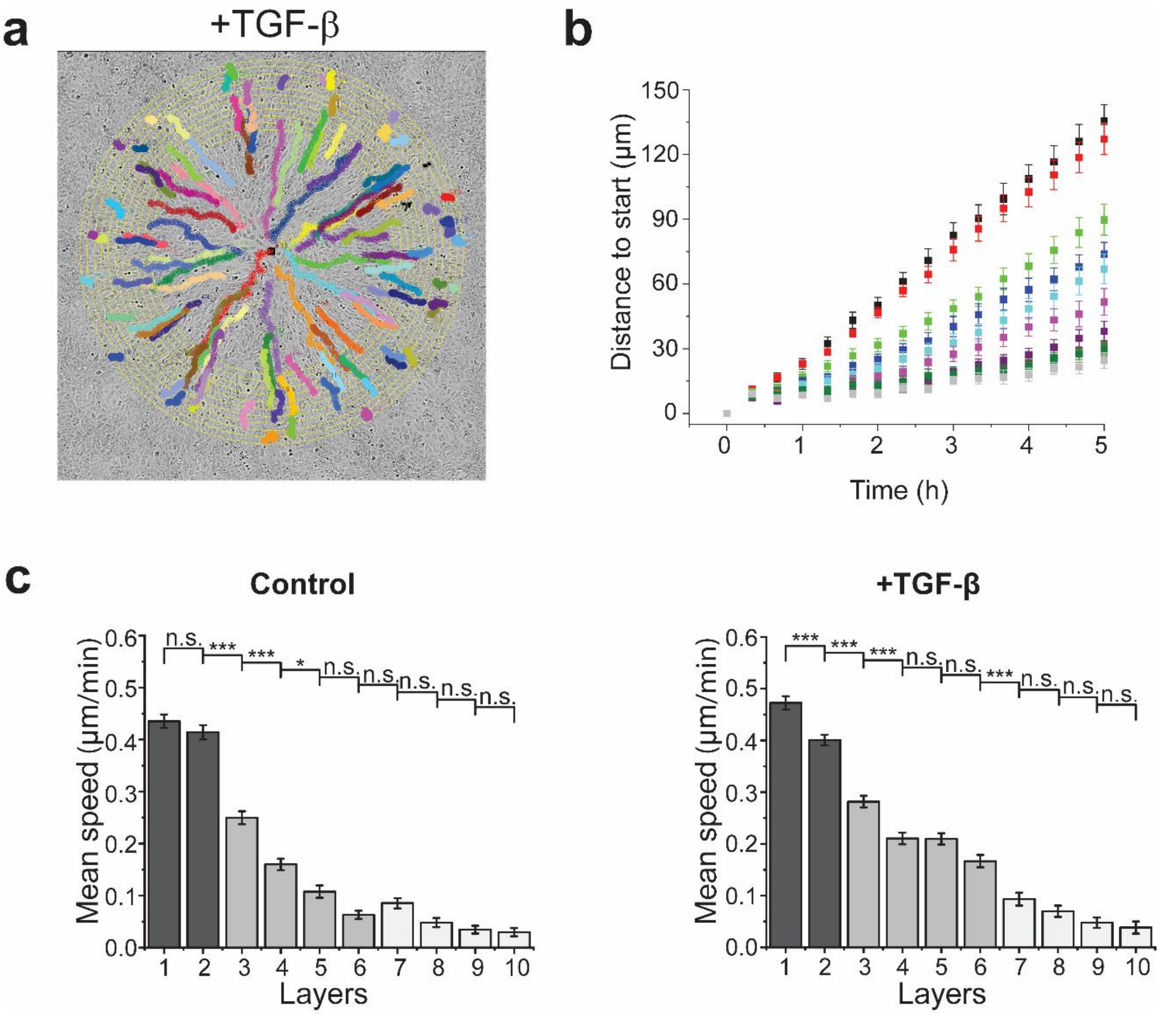


**Supplementary Fig. 4.** Migration of individual MEC1 cells during gap closure. (**a**) Trajectories of individual cells during the gap closure of MEC1 cells treated with TGF-β. Yellow circles mark the first ten layers of cells at 0 h. The thickness of each ring is set to 49 µm based on the average cell size we measured. Seven (7) cells within each layer were tracked. (**b**) Plot showing the distance to the initial positions for cells in each layer as a function of time for the first 5 hours. (**c**) Bar plots showing the mean speed of cells in each layer. Data are represented as mean ± s.e.m. n.s., $P > 0.05$, *, $P < 0.05$, ***, $P < 0.001$.

**Supplementary Figure 5**

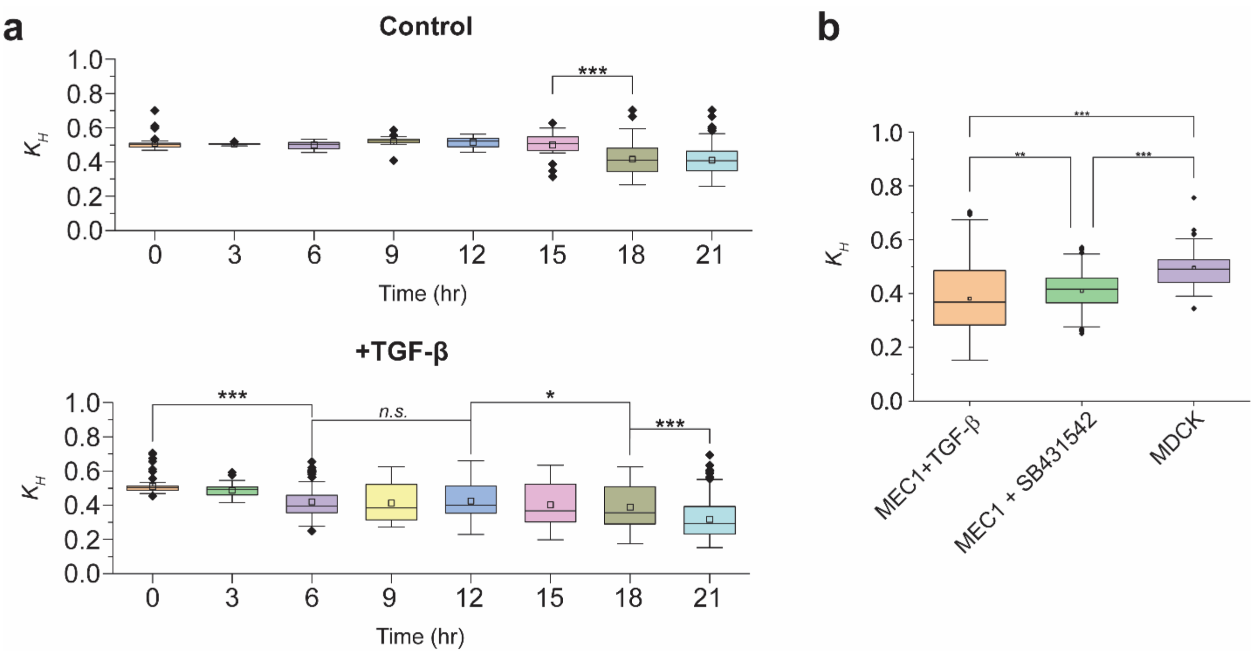


**Supplementary Fig. 5.** Orientation analysis for stress fibers in MEC1 cells. (**a**) Box plots showing $K_H$ values, which quantify the actin alignment, at different time points for control and TGF-β1 treated conditions. (**b**) Box plot showing $K_H$ values in MEC1 and MDCK cells with treatment as indicated. Data are represented as mean ± s.e.m. n.s., $P > 0.05$, *, $P < 0.05$, **, $P < 0.01$.***, $P < 0.001$.

**Supplementary Figure 6**

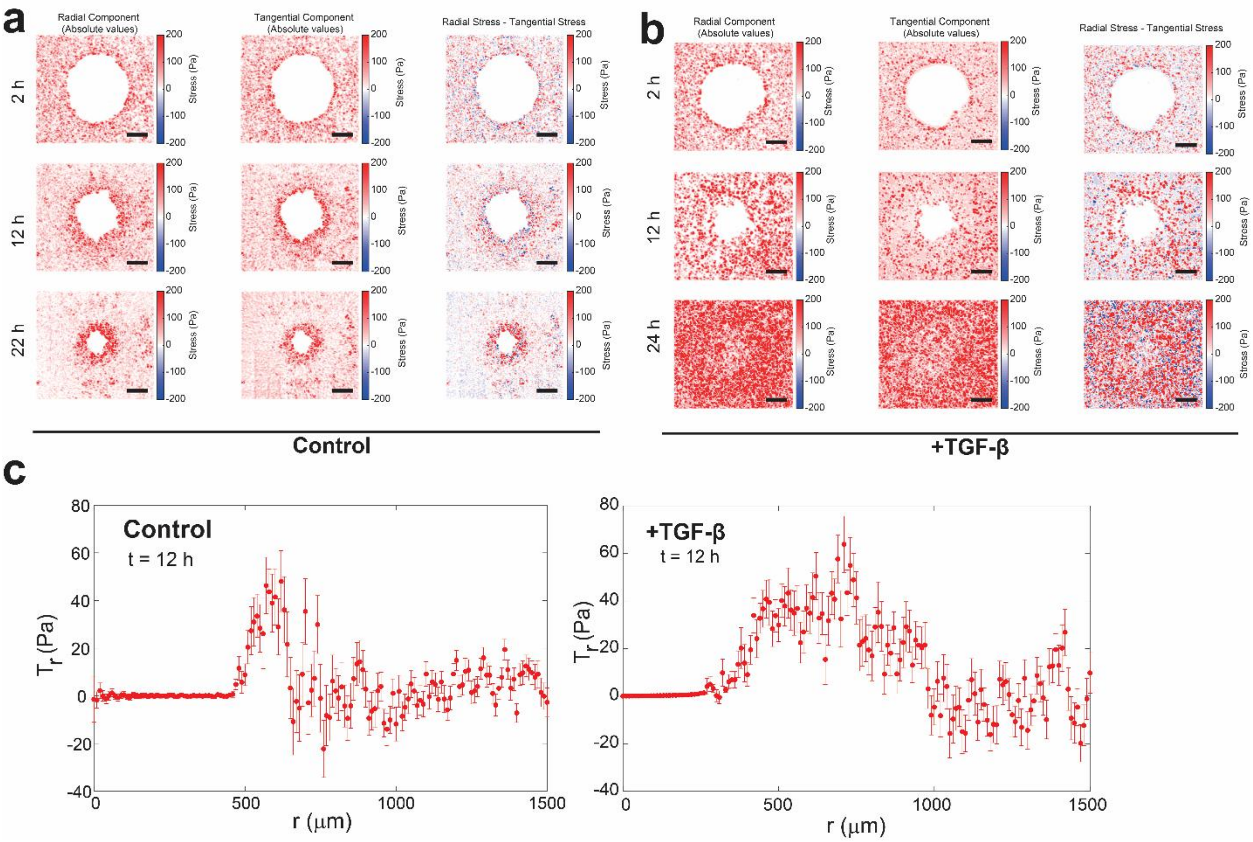


**Supplemenary Figure 6**: (**a-b**) Comparison of the magnitude of radial traction stress and tangential traction stress. Scale bar, 500 μm. (**c**) Quantification of radial traction stress as a function of distance to the gap center at 12 h.

**Supplementary Figure 7**

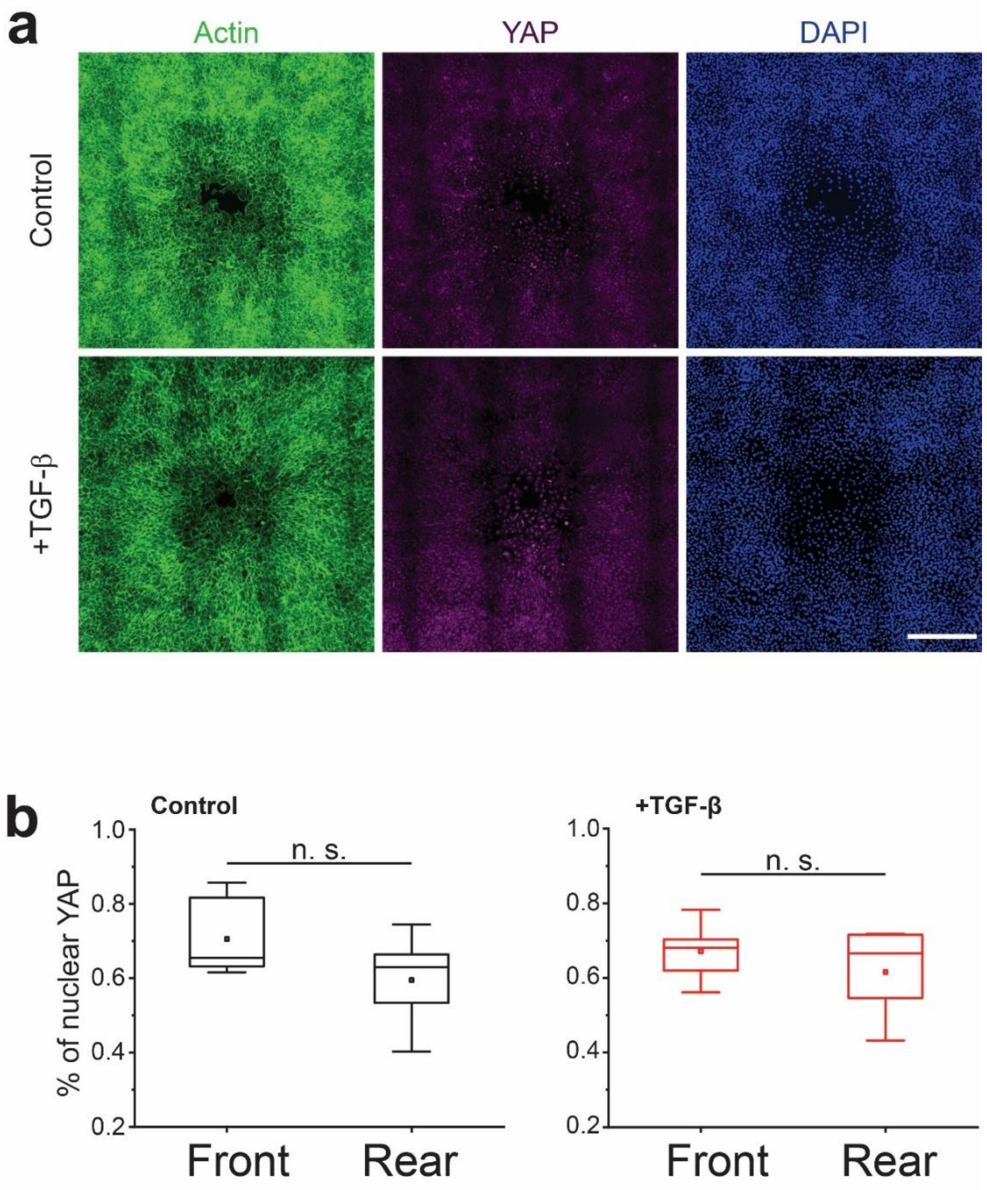


**Supplementary Fig. 7.** Enhanced contractility is YAP independent. (**a**) Representative fluorescence images showing the actin, YAP and nuclei staining in MEC1 cells at 18 h after stencil removal. (**b**) Box plot showing the percentage of cells with YAP predominantly expressed in the nuclei in the front (layer 1-2) and rear (layer 3-10). Data are represented as mean ± s.e.m. n.s., $P > 0.05$.

**Supplementary Figure 8**

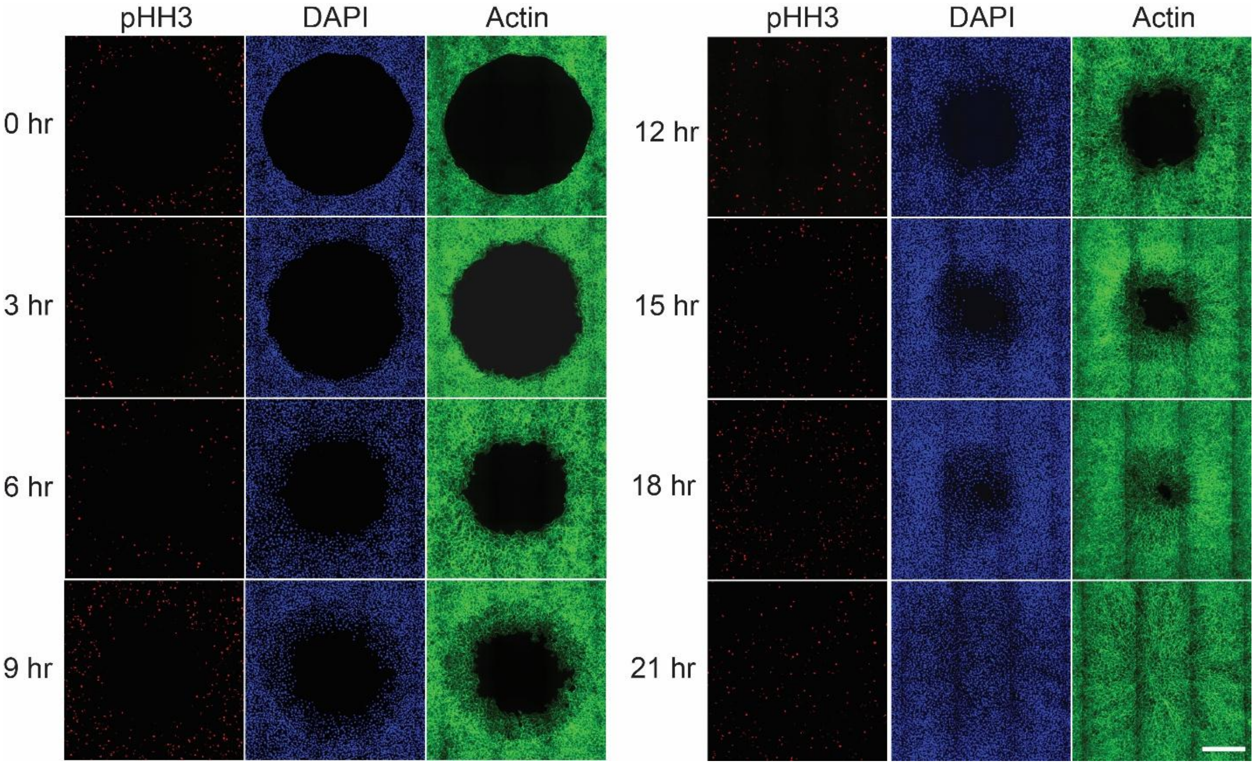


**Supplementary Fig. 8.** Representative fluorescence images showing pHH3 positive cells at different time points in MEC1 cells treated with TGF-β1. Scale bar, 500 μm.

**Supplementary Figure 9**

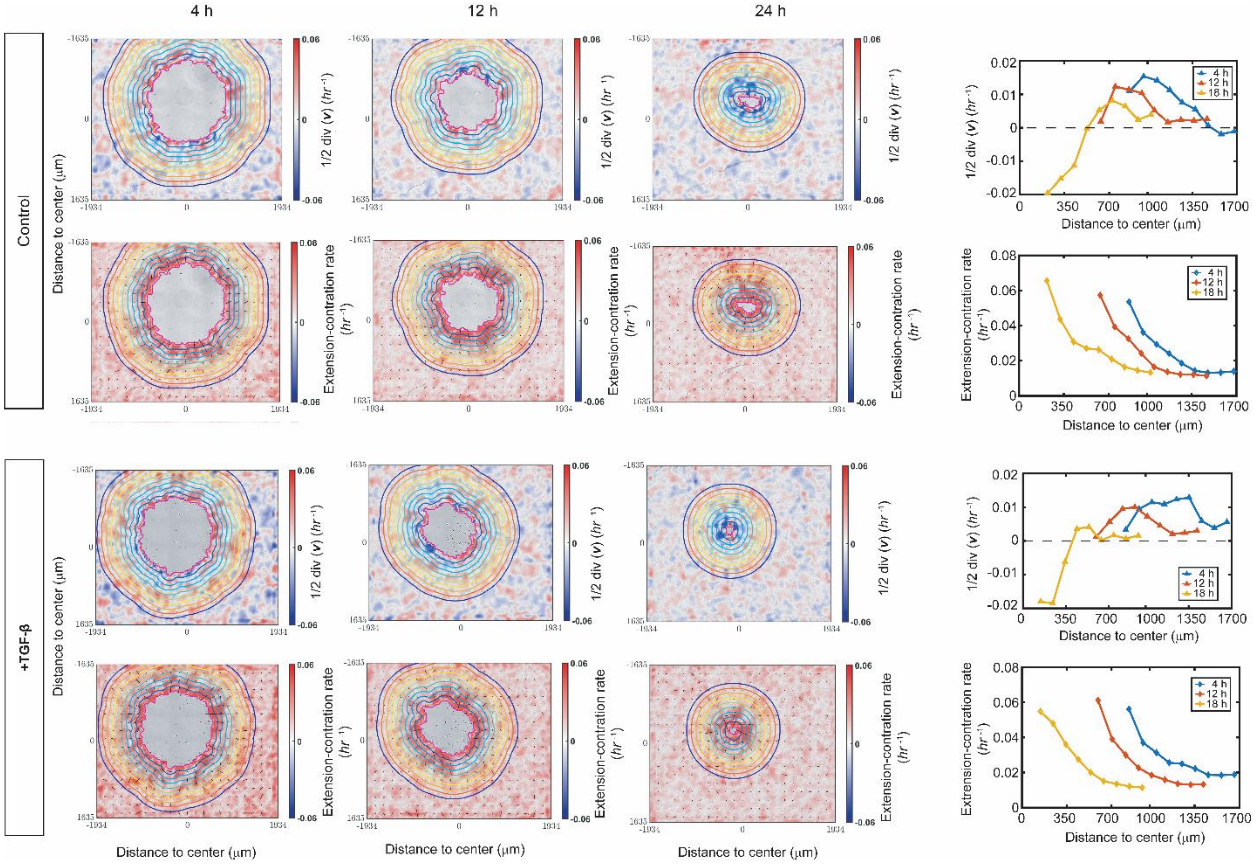


**Supplementary Fig. 9.** Two-dimensional color map of growth and extension-contraction rates at 4 h, 12 h, and 24 h. The contours divided the tissues into 9 layers and both measurements were averaged within each layer and shown in the graph to the right, which showed the distribution of growth and extension-contraction rates along the distance from the center. In the extension-contraction maps, the local direction of elongation (shown by black bars) aligned mostly with the local radial direction.

**Supplementary Figure 10**

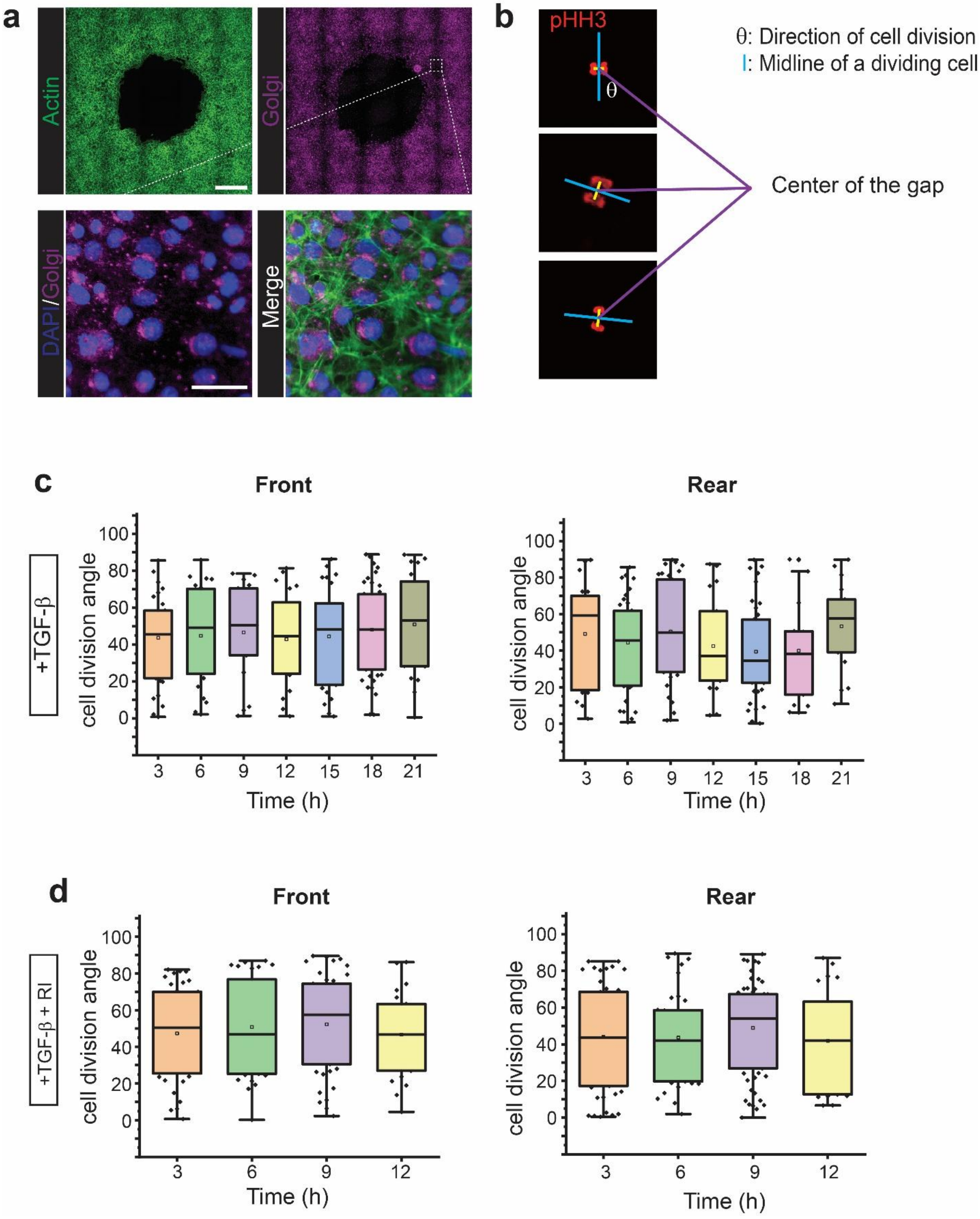

**Supplementary Fig. 10.** Oriented cell division is not involved in the tissue convergent extension. (**a**) Representative images showing the actin and Golgi apparatus in TGF-β1 treated cells at 12 h. Scale bar, 500 μm (top) and 50 μm (bottom). (**b**) Images and schematics showing pHH3 staining of a dividing cell. The direction of cell division is defined as the angle between the midline direction and the line connecting the center of the midline and the center of the gap. (**c-d**) Box plots showing cell division angle distribution throughout the gap closure process in MEC1 cells treated with TGF-β1 (c); and TGF-β1 and Y27632 (RI) (d). $n$ >18 cells at each time point. No statistical significances were found in all groups.

**Supplementary Figure 11**

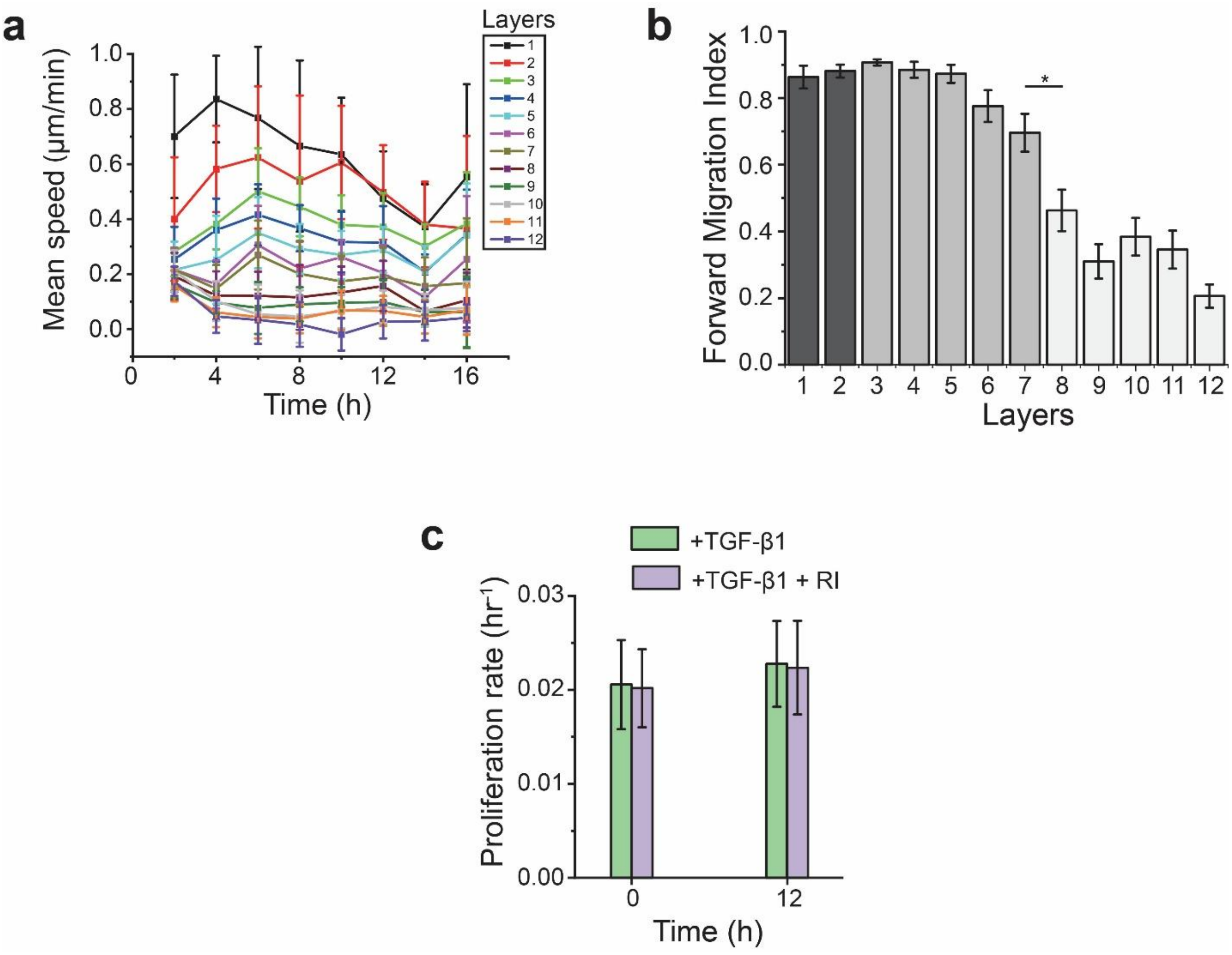


**Supplementary Fig. 11.** RI treatment increased the migration speed and directionality of MEC1 cells without affecting proliferation rate. (**a**) Plot showing the mean speed of RI treated MEC1 cells as a function of time. Twelve (12) cell layers were analyzed with a layer width of 49 μm. $n > 6$ cells in each layer. (**b**) The forward migration index of each layer in RI treated MEC1 cells. $n > 6$ cells in each layer. (**c**) Proliferation rate for MEC1 cells at 0 and 12 h during gap closure. $n = 4$ independent samples. Data are represented as mean ± s.e.m.

### Supplementary Table:

#### Table 1: summary of parameters used in the modeling

| **With fiber-reforcement** $k_H = 0.5$ | **Control**:$0 \leq \beta \leq 0.05$ **+TGF-$\boldsymbol{\beta}$**:$0 \leq \beta \leq 0.05$ **+RI**: $0.03 \leq \beta \leq 0.1$ $(\Delta\beta = 0.005)$ | $0 \leq \zeta \leq 0.05$ $(\Delta\zeta = 0.005)$ | **Control**:$10 \leq \eta \leq 100$ **+TGF-$\boldsymbol{\beta}$**: $10 \leq \eta \leq 100$ $(\Delta\eta = 10)$ **+RI**: $5 \leq \eta \leq 20,\ (\Delta\eta = 5)$ | **Control**:$0.012 \leq \gamma \leq 0.024$ **+TGF-$\boldsymbol{\beta}$**:$0.02 \leq \gamma \leq 0.028$ **+RI**: $0.022 \leq \gamma \leq 0.032$ $\Delta\gamma = 0.002$ |
|---|---|---|---|---|
| **Control** | 0.025 | 0.015 | 20 | 0.02 |
| **+TGF-$\boldsymbol{\beta}$** | 0.01 | 0.01 | 20 | 0.024 |
| **+RI** | 0.06 | 0.02 | 15 | 0.026 |
